# A synthetic PGC1α co-activator reprograms microglial immunometabolism to preserve cone function in retinal degeneration

**DOI:** 10.64898/2026.09.29.753881

**Authors:** Thomas Winogrodzki, Jayvin Hernandez-Reyes, Rajvir Solanky, Richard Lowery, Georgy Komissarov, Wen Huang, Boyuan Wang, Neoklis Makrides, Sharifa Davis, Siyuan Liu, Yong-Shi Li, Wen-Hsuan Wu, Nicholas Nolan, Robert van de Werken, Aykut Demirkol, Anders Knudsen, Nuntachai Surawatsatien, Yao Li, James Hurley, Stephen H Tsang

## Abstract

Retinitis pigmentosa (RP) is genetically heterogeneous, yet initiating mutations converge on primary rod degeneration followed by cone dysfunction and loss. Activated microglia are implicated in this secondary phase, but whether selective immunometabolic reprogramming of microglia preserves retinal function remains unknown. In human iPSC-derived microglia and retinal CD11b⁺ cells, PGC1α induction suppressed LPS-driven interferon signaling and cytokine release, yet Seahorse analysis and isotopologue tracing showed that energy metabolism remained altered. We therefore fused PGC1α to a multivalent p65-HSF1 module, yielding ∼2.5-fold higher nuclear receptor output than native PGC1α and coordinated engagement of oxidative and anti-inflammatory programs in mouse retinal CD11b⁺ cells. Microglia-restricted expression in humanized *RHO^P347L^* mice preserved photopic responses through six months, increased retinal thickness and promoted ramified subretinal IBA1⁺ morphology. Microglial immunometabolism thus represents a potentially genotype-independent neuroprotective target in RP, and engineered PGC1α co-activators provide a means to sustain oxidative metabolism under inflammatory pressure.

---

Retinitis pigmentosa (RP) globally affects approximately 1.5 million people and causes progressive rod loss followed by secondary cone degeneration and severe visual impairment ^1^. Pathogenic variants in more than 80 genes span autosomal-dominant, autosomal-recessive and X-linked disease ^2^. Gene replacement and editing benefit selected monogenic subtypes ^3,4^ but remain mutation-restricted and may provoke retinal inflammation ^5^, motivating gene-agnostic strategies directed at shared disease mechanisms.

Neuroinflammation actively contributes to RP progression. Sustained cytokine, chemokine and oxidative-stress signals occur in patient ocular fluids across disease stages ^6^. This axis is pharmacologically accessible in humans: interleukin-6 receptor blockade reduced cystoid macular edema in inflammation-predominant RP ^7^, whereas minocycline improved cone-driven flicker electroretinogram amplitudes in an open-label trial ^8^. Neither intervention is durable or cell-selective, and neither directly targets the metabolic state that sets inflammatory output in tissue-resident immune cells.

Retinal degeneration activates and redistributes resident microglia, while inflammatory activation drives glycolytic reprogramming that sustains cytokine output ^9,10^. Microglial depletion limits photoreceptor loss in rd10 mice ^11^, and interventions targeting TREM2 ^12^, CCL7-STAT1 ^13^, TGF-β1 ^14^ or RPE-derived PD-L1 ^15^ also reduce degeneration or preserve cones. These studies establish microglial state as therapeutically consequential, but do not test whether durable, cell-restricted immunometabolic reprogramming alone can preserve cone function.

Peroxisome proliferator-activated receptor gamma co-activator-1 alpha (PGC1α) is a transcriptional co-activator of mitochondrial biogenesis and oxidative metabolism that can restrain inflammatory programs ^16^. Monocyte-lineage PGC1α elevation via Cx3cr1-Cre limits tissue injury and cytokine output in acute ischemic brain injury ^17^, but chronic neurodegenerative contexts and microglial specificity remain unaddressed. PGC1α has been implicated in retinal degeneration itself: the PGC1α/NRF-1/TFAM mitochondrial-biogenesis axis is downregulated in RP mouse models, and its pharmacological restoration with the PGC1α agonist ZLN005 delays photoreceptor loss ^18^, while combined PPARα, PPARγ, and PGC1α activation reduces photoreceptor death in retinal explants ^19^. However, these studies have not addressed PGC1α’s contribution within retinal microglia, and most microglial immunometabolic work has relied on brain tissue or BV-2 cell lines, which diverge transcriptionally and metabolically from retinal microglia ^20,21^. Whether retinal microglial PGC1α reprogramming alone can reduce inflammation and protect photoreceptors remains unresolved.

We addressed this in three steps. We first tested ZLN005 in human iPSC-derived microglia and donor-retina CD11b⁺ cells. Because inflammation suppresses endogenous PGC1α activity ^16,22^, we then fused *PGC1α* to a multivalent *p65-HSF1* module inspired by modular CRISPR activators ^23^, creating Activ_PGC1α to increase output per molecule. Finally, we tested microglia-restricted Activ_PGC1α in our humanized hu*RHO^P347L^* model ^24,25^, which carries the severe early-onset human *RHO P347L* allele.

## Results

### PGC1α agonist ZLN005 reduces microglial LPS-mediated inflammatory response but does not prevent LPS-associated glycolytic shift in iPSC-derived microglia

Based on the finding that intravitreal PGC1α agonist ZLN005 rescues photoreceptor function in a mouse RP model ^18^, we asked whether retinal microglia are among the main effectors. We compared bulk transcriptional profiling of treated maturated human iMGs with 20 µM ZLN005 for 48 h and/or 1 µg/ml LPS for the final 6 h to untreated controls (**Fig. 1A**).

**Fig. 1.**
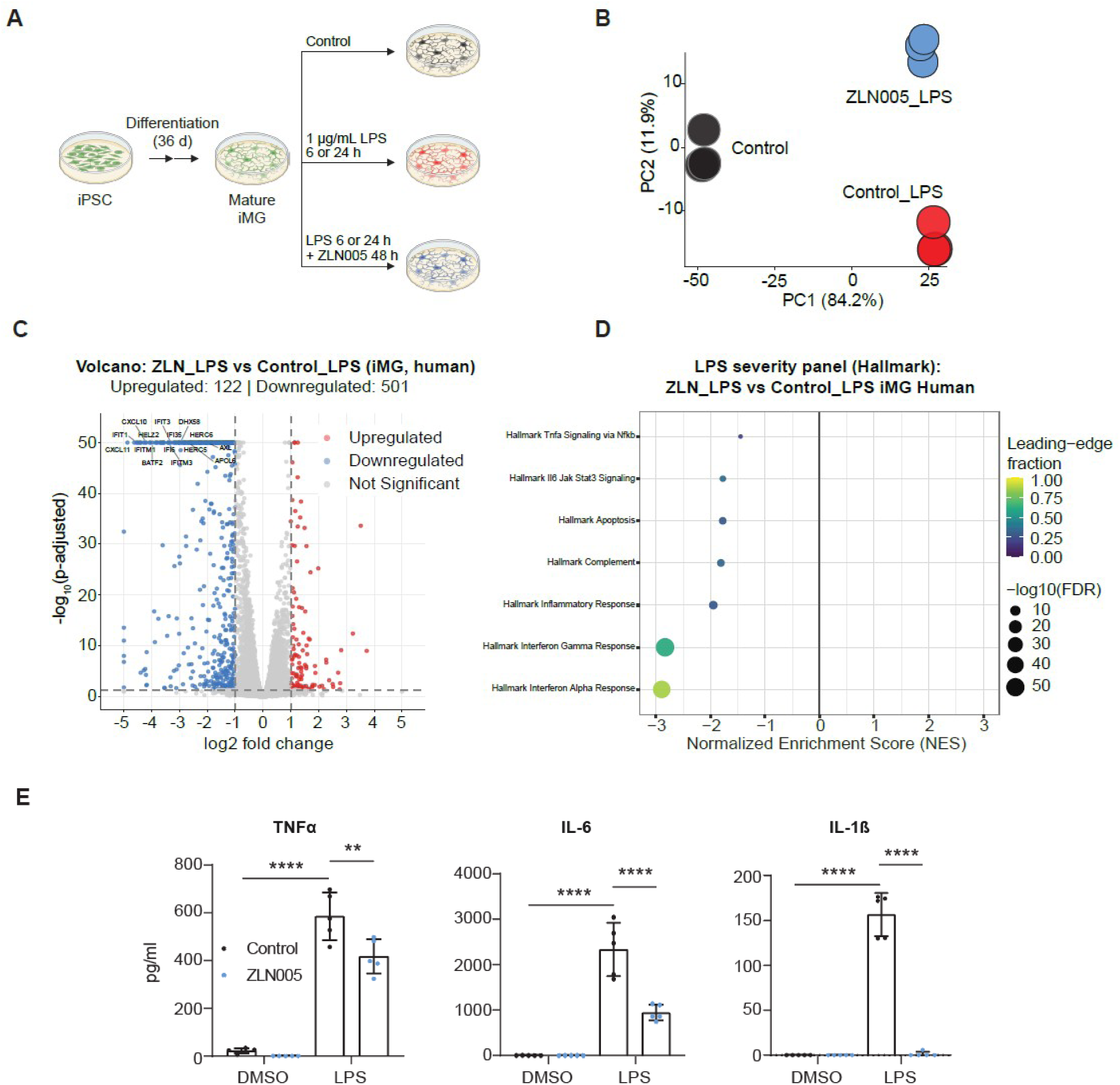
PGC1α agonist ZLN005 dampens the microglial LPS inflammatory program in human iPSC-derived microglia (iMG). (**A**) Experimental design: human iPSCs were differentiated to mature iMG (36 days) and treated with vehicle (control), 1 µg/ml LPS, or 20 µM ZLN005 (48 h) with LPS added for the final 6 h, followed by bulk RNA-sequencing. (**B**) Principal component analysis (PCA) of iMG transcriptomes; LPS drives separation along PC1 (84.2% of variance) and ZLN005 co-treatment along PC2 (11.9%). (**C**) Volcano plot of the ZLN005+LPS versus control+LPS contrast (122 upregulated, 501 downregulated genes; DESeq2 Wald test, adjusted *P* < 0.05). (**D**) Targeted Hallmark gene-set enrichment for seven inflammatory pathways, all showing significant negative enrichment under ZLN005 co-treatment; interferon alpha and gamma responses are most strongly suppressed. NES, normalized enrichment score. (**E**) Multiplexed immunoassay of secreted TNFα, IL-6 and IL-1β in culture supernatants (control [DMSO] versus LPS, with or without ZLN005). n = 5 each, values below the lower limit of detection were set to 0.

Principal component analysis (PCA) revealed clear separation of the three conditions (**Fig. 1B**). Untreated controls and both LPS-stimulated groups segregated predominantly along PC1 (84.2% of variation), indicating that LPS exposure constitutes the dominant source of transcriptomic variation. ZLN005-treated LPS samples separated from the LPS-only condition primarily along PC2 (11.9%), suggesting that ZLN005 imposes a distinct transcriptional signature on the inflammatory state rather than reverting it fully to baseline. Unsupervised hierarchical clustering of the top 500 most variable genes corroborated this pattern (**Supplementary Fig. 1A, B**). Differential expression analysis of ZLN005_LPS versus Control_LPS identified 623 differentially expressed genes (122 upregulated, 501 downregulated), with the top 15 most significant genes all downregulated interferon-response genes (*APOL6, AXL, BATF2, CXCL10, CXCL11, DHX58, HELZ2, HERC5, HERC6, IFI35, IFI6, IFIT1, IFIT3, IFITM1, IFITM3*), reflecting a coordinated, near-complete suppression of the type-I interferon signature (**Fig. 1C**). Gene set enrichment analysis (GSEA) against the Reactome database confirmed that the downregulated arm was dominated by interferon alpha/beta signaling, interferon gamma signaling, and antiviral modules including ISG15-mediated antiviral mechanisms and DDX58/IFIH1-mediated induction of interferon alpha/beta (**Supplementary Fig. 1C**). The comparatively few positively enriched sets were not directly linked to inflammation, reflecting rRNA modification and processing, the RND2 GTPase cycle, and cilium assembly. A targeted analysis of seven predefined Hallmark inflammatory pathways showed significant negative enrichment for all seven, with interferon alpha and gamma responses most strongly suppressed, followed by inflammatory response, complement, apoptosis, IL6-JAK-STAT3, and TNFα signaling via NF-κB (**Fig. 1D**).

At the gene level, we examined transcripts spanning inflammatory signaling and mitochondrial biogenesis across the three contrasts (**Supplementary Fig. 2**). Boxplot analysis of 25 representative transcripts showed that interferon-stimulated genes, JAK-STAT components, and inflammatory chemokines were uniformly LPS-induced and significantly attenuated by ZLN005 co-treatment, whereas mitochondrial biogenesis transcripts showed a mixed response. The cytosolic DNA sensor *CGAS*, but not *STING1*, was rescued by ZLN005.

Validation at the protein level using multiplexed immunoassays of culture supernatants collected after 24 h of LPS confirmed significantly reduced secreted TNFα, IL-6, and IL-1β in ZLN005+LPS relative to LPS alone (**Fig. 1E**). This reduction of pro-inflammatory cytokines indicates a shift of microglial phenotypes toward a less inflammatory state.

We next asked whether this de-escalation was accompanied by the oxidative metabolic shift classically associated with PGC1α ^17^. Seahorse real-time ATP-rate assays of cells pre-treated with LPS and/or ZLN005 or controls resolved the underlying bioenergetics (**Fig. 2A, B; S3A, B**). Under homeostatic conditions, ZLN005 produced the expected oxidative phenotype with significantly elevated basal oxygen consumption and mitochondrial ATP production, placing cells in the oxidative quadrant of the metabolic phenotype map. Contrary to expectation, however, this oxidative polarization was abolished under LPS co-stimulation. Mitochondrial ATP production fell below vehicle control while basal extracellular acidification increased, placing ZLN005+LPS cells in the highly glycolytic quadrant adjacent to LPS-only cells, which we termed metabolic override. In line, all three perturbations elevated extracellular lactate relative to vehicle (control 600 µM; ZLN005 alone 940 µM; LPS alone 860 µM; ZLN005+LPS 1030 µM), with the combination significantly exceeding LPS alone (**Fig. 2C**).

**Fig. 2.**
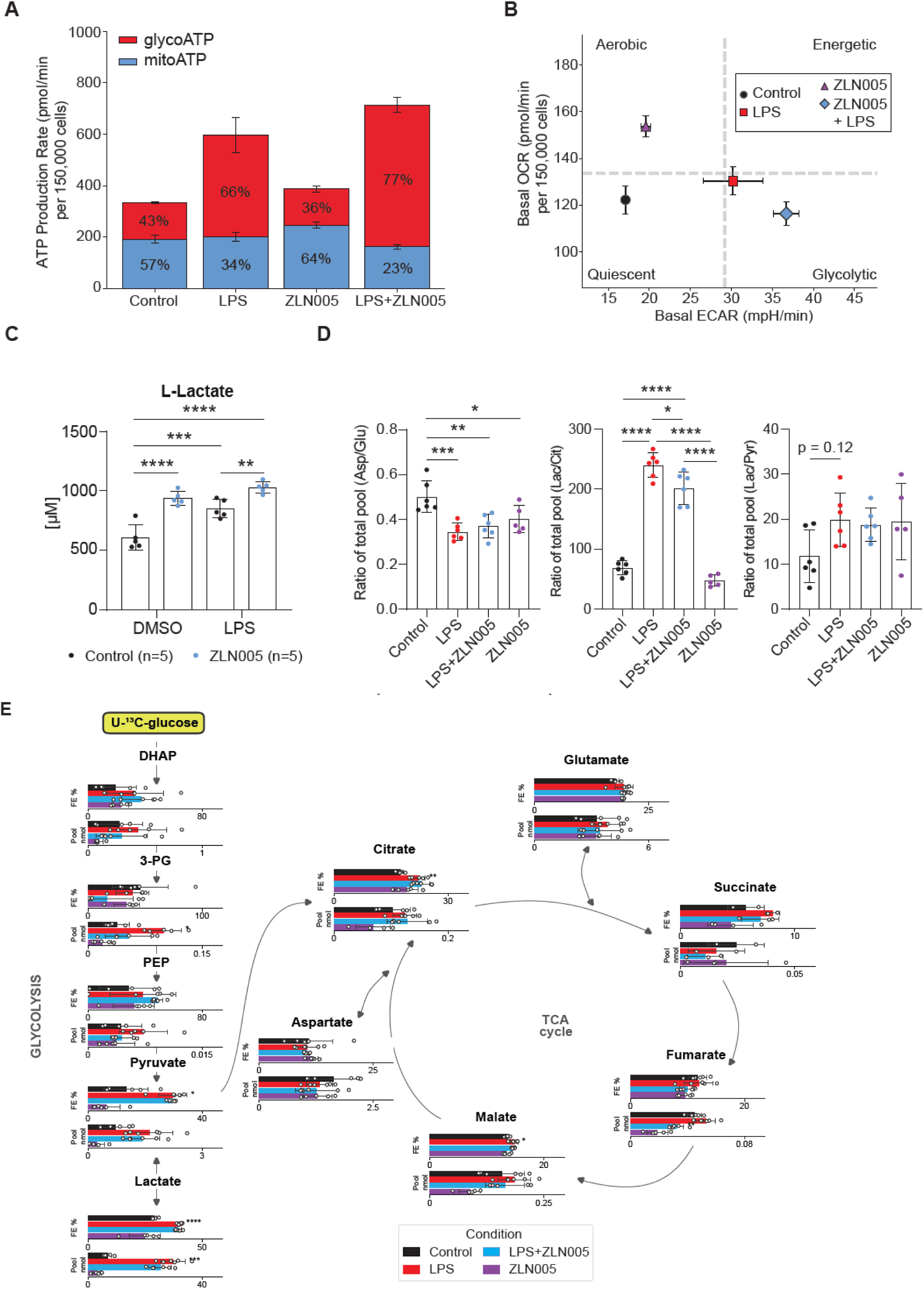
Inflammatory load overrides the ZLN005-induced oxidative program in iPSC-derived microglia (iMG). (**A**) Seahorse ATP-rate assay showing mitochondrial (oxidative, mitoATP) and glycolytic (glycoATP) ATP-production fractions. (**B**) OCR-ECAR map: ZLN005 shifts homeostatic cells toward the oxidative quadrant, whereas ZLN005+LPS cells occupy the glycolytic quadrant beside LPS-only cells ("metabolic override"). (**C**) Extracellular L-lactate (µM), significantly higher with ZLN005+LPS than LPS alone. (**D**) ¹sC-glucose total-pool ratios: aspartate/glutamate (Asp/Glu), lactate/citrate (Lac/Cit) and lactate/pyruvate (Lac/Pyr). With inflammation, Lac/Cit rises and remains elevated with LPS+ZLN005 but is lower than with LPS alone; Asp/Glu falls; Lac/Pyr trends nonsignificantly (P = 0.12). (**E**) [U-¹sC]glucose tracing through glycolysis and the TCA cycle. Each node shows fractional ¹sC enrichment (FE%, upper) and total pool (Pool nmol, lower) as horizontal bars for Control (black), LPS (red), LPS+ZLN005 (blue) and ZLN005 (purple), with biological replicates as points; grey arrows show carbon flow. iMG (150,000 cells/well) were matured with 20 mM ¹²C-glucose + 100 µM ¹²C fatty acid-BSA; ZLN005 (20 µM) was present for 48 h and LPS (1 µg/ml) for the final 24 h. Tracing used a 6 h pulse of 20 mM [U-¹sC]glucose + 100 µM ¹²C fatty acid-BSA with LPS and ZLN005 maintained. Metabolites were analyzed by GC-MS. OCR, oxygen consumption rate; ECAR, extracellular acidification rate. Data are mean ± SEM (A, B) or s.d. (C-E); n = 6 (ZLN005, LPS+ZLN005) and n = 3 (Control, LPS) for Seahorse; n = 6 (Control, LPS, LPS+ZLN005) and n = 5 (ZLN005 alone) for isotopologue tracing. *For (D), *P < 0.05, **P < 0.01, ***P < 0.001, ****P < 0.0001. For (E), *q < 0.05, **q < 0.01, ***q < 0.001, ****q < 0.0001*. Ratios in (D) were compared by one-way ANOVA with Tukey’s test; node-level enrichment and pool sizes in (E) by two-sample t-tests with Benjamini-Hochberg FDR correction across all node-level tests (q).

To resolve substrate routing directly, we traced U-¹sC-glucose through glycolysis and the TCA cycle by stable isotope-resolved GC-MS and summarized the resulting shifts as total-metabolite pool ratios (**Fig. 2D, E**). The lactate-to-citrate ratio, an index of glycolytic overflow relative to oxidative TCA engagement, rose sharply with inflammation (Control 69 ± 12 to LPS 240 ± 21; one-way ANOVA, P < 0.0001) and behaved as a quantitative readout of the metabolic override: LPS+ZLN005 (202 ± 27) was reduced relative to LPS (p = 0.01) yet remained far above ZLN005 alone (47 ± 9), which itself numerically fell below control, consistent with an oxidative shift under homeostasis that is erased by TLR4 signaling. The aspartate-to-glutamate ratio decreased under inflammation (Control 0.50 ± 0.07 to LPS 0.35 ± 0.04; P < 0.001) and was not restored by ZLN005. Of note, aspartate fell relative to every other measured pool (Cit/Asp, Glu/Asp, Mal/Asp, 3PG/Asp and Pyr/Asp all q < 0.01). Transcriptionally, this reflects simultaneous restriction of synthesis and increased consumption, as LPS downregulated both aspartate aminotransferases (*GOT1*, *GOT2*; padj < 10⁻⁶) while strongly inducing Argininosuccinate-Synthetase 1 (*ASS1*; ≈55-fold, padj < 10⁻²⁷⁰) and asparagine synthetase (*ASNS*; padj = 1.6 × 10⁻s), neither of which was rescued by ZLN005 (**Supplementary Fig. 2**). Aspartate depletion therefore occurred despite increased pyruvate-carboxylase-dependent replenishment of oxaloacetate (M3/M2 aspartate 0.28 to 0.43; P < 0.0001), consistent with cataplerotic demand rather than with a respiratory constraint on aspartate synthesis. The lactate-to-pyruvate ratio trended higher with activation but did not reach significance (Control 11.8 ± 5.8 to LPS 19.9 ± 5.9; P = 0.12), indicating that a change in the cell’s redox state was likely not the driving factor of the metabolic override.

LPS markedly increased glucose incorporation into lactate (M3 lactate enrichment 29 to 40%, q < 10⁻⁶; total lactate pool ∼4-fold expanded) and pyruvate (M3 pyruvate 13 to 30%, q = 0.016), together with a smaller increase in labeling of the TCA intermediates citrate and malate (M2 citrate 17 to 22%, q = 0.006; M2 malate q = 0.021), indicating that inflammatory activation raises overall glucose utilization while committing most carbon to lactate. Under homeostatic conditions, ZLN005 alone contracted glycolytic and TCA pools to 0.32-0.64× of control while fractional labeling of the TCA segment was preserved or selectively increased, with higher M2-glutamate (17.7 to 19.4%, q = 0.006) and M2-and M3-aspartate enrichment and a concordant trend for M2-citrate, indicating a glucose-sparing oxidative state. Critically, this oxidative routing was lost under inflammatory load. Adding ZLN005 to LPS did not shift glucose partitioning significantly compared to LPS alone at any node (LPS+ZLN005 vs LPS, all comparisons non-significant), and the combination resembled LPS rather than ZLN005 alone (M3 lactate, M3 pyruvate, and M2 malate all q < 0.05 versus ZLN005). Only the significant reduction in lactate/citrate ratio (p = 0.010) indicates partial reduction in LPS-mediated metabolic shifts. Thus, at the level of carbon flux, ZLN005 imposes a modest oxidative program under homeostatic conditions that is mostly overridden by concurrent TLR4 signaling, directly corroborating the respirometry and lactate data.

Crucially, this loss of metabolic reprogramming did not abolish the anti-inflammatory effect. ZLN005 strongly suppressed LPS-induced cytokine secretion even as mitochondrial ATP production declined and glycolytic carbon accumulation intensified, suggesting that the anti-inflammatory action remains engaged even when the inflamed metabolome cannot be forced back toward oxidative phosphorylation. However, whether the metabolic override presents a limitation to the anti-inflammatory potential of native PGC1α is unclear.

### ZLN005 pre-conditioning reshapes the microglial secretome toward a less inflammatory paracrine signal

The lower intracellular lactate pool under ZLN005 despite increased extracellular lactate accumulation (**Fig. 2C, E**) suggests faster lactate turnover, without establishing reduced glycolytic production. To determine whether ZLN005 pre-conditioning changes the net pro-inflammatory inductive potential of the microglial secretome despite persistent lactate export, we performed a two-stage conditioned-medium transfer assay (**Supplementary Fig. 4A**). iMGs were treated with ZLN005 and/or LPS as before, washed thoroughly to remove residual ZLN005, and re-challenged with LPS alone before collecting donor supernatants. Independently differentiated, naive recipient iMG exposed to the ZLN005-conditioned secretome showed significantly reduced IL6 induction (∼900-fold versus ∼2000-fold; p < 0.05) and a trend toward reduced TNFα (∼7-fold versus ∼12-fold; p = 0.07) (**Supplementary Fig. 4B**). This indicates that ZLN005 reshapes the paracrine signal to carry reduced pro-inflammatory instruction despite metabolic override.

### PGC1α agonism suppresses the LPS inflammatory program in primary human donor-eye-derived retinal microglia

To test whether these findings extend to primary human retinal cells, we isolated rMGs from three human post-mortem donor eyes (female donors aged 29, 51, and 54 years; **Supplementary Table 1**) and profiled transcriptomes of FACS-sorted CD11b^+^ cells treated with LPS or LPS+ZLN005 (**Supplementary Fig. 5A; Fig. 3A, B**). As expected for primary human material, PCA clustering was dominated by inter-donor variation, and the donor-paired ZLN005 effect was comparatively subtle (**Fig. 3C**). The LPS+ZLN005 versus LPS contrast in donor-blocking design yielded 54 significant differentially expressed genes (23 up, 31 down) (**Fig. 3D**). Because RNA input was low, and thus individual extreme-fold change estimates emerged, interpretation was anchored on pathway-level enrichment and directionally coherent gene modules (**Supplementary Fig. 5A, B**). ZLN005 co-treatment consistently and significantly dampened innate immune activation, protein synthesis, and extracellular-matrix remodeling, while several anabolic and growth-promoting pathways trended towards induction without reaching significance (**Supplementary Fig. 5C**). The targeted Hallmark analysis recapitulated the iMG results: all seven inflammatory pathways showed negative enrichment, with interferon alpha and gamma responses again most strongly suppressed, followed by inflammatory response, IL6-JAK-STAT3 signaling, and complement, with apoptosis the weakest and non-significant arm (**Fig. 3E**). Across the human iMG and donor rMG datasets, the same core interferon-centered program was suppressed, establishing that PGC1α agonism drives reproducible anti-inflammatory reprogramming and positioning retinal microglia as plausible key effectors of ZLN005-directed neuroprotection. The absence of upregulated oxidative pathways in both systems confirms that the inflammatory override is not an artifact of iPSC-derived cells.

**Fig. 3.**
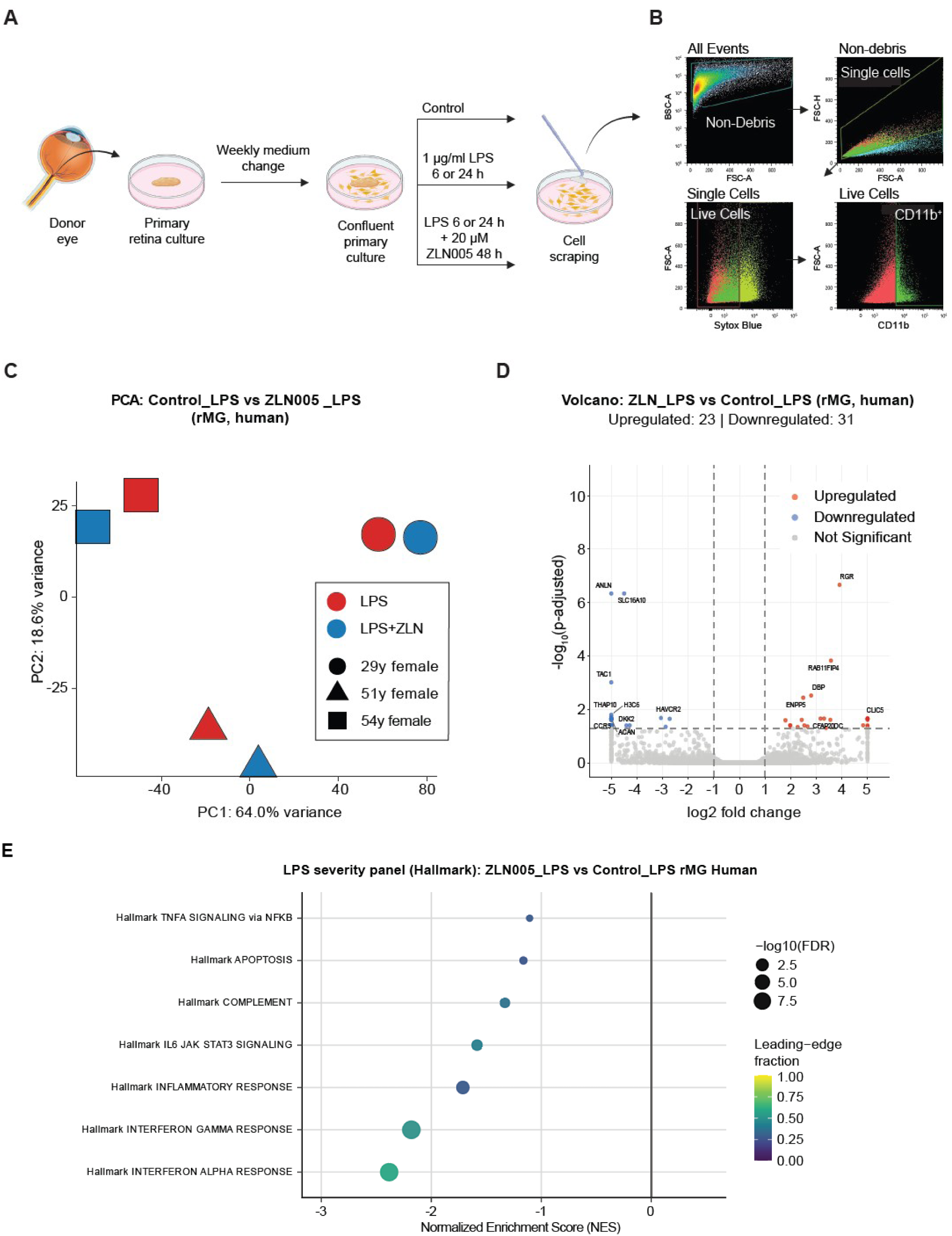
PGC1α agonism suppressed the LPS inflammatory program in primary human donor-eye retinal microglia (rMG). (**A**) Workflow: retinal microglia were isolated from post-mortem human donor eyes, cultured, treated with LPS or LPS+ZLN005, and processed for transcriptomics. (**B**) FACS gating strategy for isolation of CD11b+ retinal microglia (sequential gating on non-debris, single, live, CD11b^+^ cells). (**C**) PCA of rMG transcriptomes from three donors (female, aged 29, 51 and 54 years); variation is dominated by inter-donor differences (PC1 64.0%, PC2 18.6%). (**D**) Volcano plot of the LPS+ZLN005 versus LPS contrast in a donor-blocked design (23 upregulated, 31 downregulated genes; DESeq2 Wald test, adjusted *P* < 0.05). (**E**) Targeted Hallmark enrichment for seven inflammatory pathways, all negatively enriched under ZLN005 co-treatment, recapitulating the iMG result with interferon alpha and gamma responses most strongly suppressed. NES, normalized enrichment score.

### Engineering Activ_PGC1α to bypass the metabolic override

Because active immune responses are known suppressors of PGC1α activity ^16,22^, which might permit metabolic override, we sought to raise the intrinsic per-molecule output of the co-activator so that pathway engagement could be sustained against an inflammatory background. Fusing transactivation domains (TAD) to nuclear effectors amplifies transcriptional output, most prominently in CRISPR-based activation systems ^23,26^. We hypothesized that applying this logic to a transcriptional co-activator rather than a programmable DNA-binding scaffold would preserve the native partner-receptor selectivity of PGC1α while increasing downstream activation amplitude, confining hyperactivation to PGC1α-responsive nuclear receptors. A trimerized p65-HSF1 TAD, termed Activ, was shown to induce a ten-fold increase in activation efficiency compared to VPR while driving substantial gene activation across ten endogenous target genes ^26^. We therefore fused the Activ-TAD to PGC1α to generate the synthetic Activ_PGC1α transcriptional co-activator, a generalizable strategy we termed TUNE (**T**AD-mediated **U**pregulation of **N**uclear Receptor **E**ngagement) (**Fig. 4A**).

**Fig. 4.**
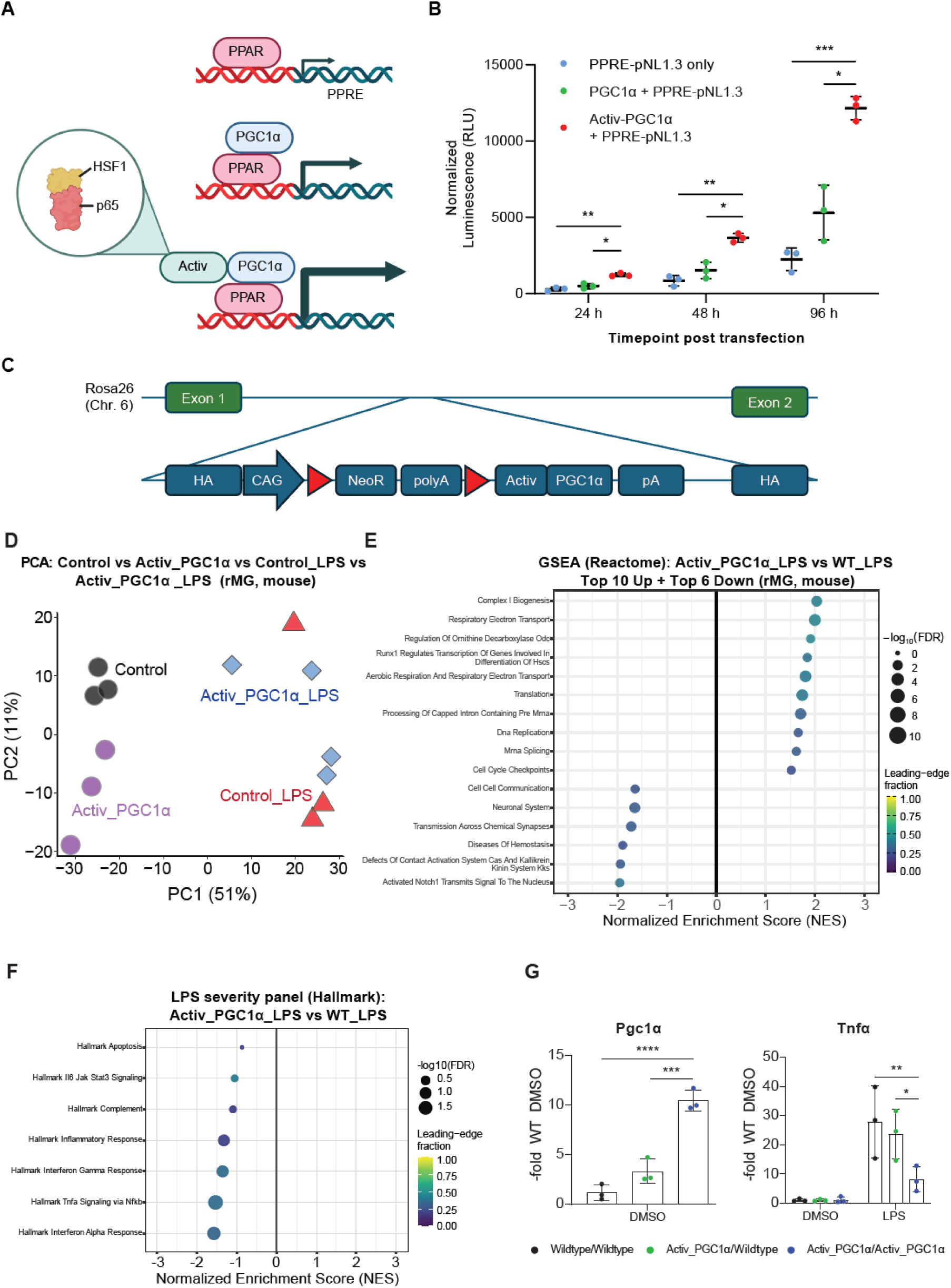
Engineering and validation of Activ_PGC1α (TUNE), a synthetic high-output PGC1α co-activator, in primary mouse retinal microglia (rMG). (**A**) Design of the TUNE co-activator: the trimerized p65-HSF1 transactivation domain (Activ) is fused to PGC1α (Activ_PGC1α), which engages nuclear-receptor partners at PPAR/PPRE elements. (**B**) PPRE-driven secreted NanoLuc luciferase reporter activity in HEK cells expressing Activ_PGC1α versus native PGC1α (normalized to P2A-co-expressed eGFP) at 24, 48 and 96 h, showing durably increased co-activator output (248%, 242% and 230% of native PGC1α). (**C**) *Rosa26* targeting strategy for the Cre-inducible *R26^LSL-Activ_PGC1α^* allele. (**D**) PCA of primary mouse rMG transcriptomes (wildtype versus tamoxifen-induced Activ_PGC1α, ± LPS); LPS is the major axis of variation (PC1 51%, PC2 11%). (**E**) Reactome GSEA (Activ_PGC1α+LPS versus wild-type+LPS): the most strongly induced pathways are mitochondrial/oxidative (Complex I biogenesis, respiratory electron transport, aerobic respiration). (**F**) Targeted Hallmark enrichment for seven inflammatory pathways, showing negative enrichment across all seven with significant suppression of the interferon alpha response and TNFα signaling via NF-κB. (**G**) qPCR showing a gene-dosage-dependent increase in *Pgc1α* transcript (up to ∼10-fold over wild type) and a corresponding gene-dosage-dependent reduction in LPS-induced *Tnfα* (wild-type → heterozygous → homozygous co-activator microglia). NES, normalized enrichment score. Data are mean ± s.d.; statistical details are given in the Methods.

Using a PPARγ-responsive element-secreted NanoLuc luciferase reporter ^27^, we found that *Activ_PGC1α* expression in HEK cells increased luminescence to 248% (24 h), 242% (48 h), and 230% (96 h) of native PGC1α, normalized to co-expressed eGFP, confirming that the Activ domain durably boosts PGC1α co-activator function (**Fig. 4B**).

### *Activ_PGC1α* expression couples anti-inflammatory and mitochondrial-oxidative programs in primary mouse microglia

We generated Cre-inducible *R26^LSL-Activ_PGC1α^* mice and crossed them to *Tmem119^CreERT^*^2^ mice for microglia-restricted, tamoxifen-inducible expression of *Activ_PGC1α* (**Fig. 4C**). CD11b^+^ primary rMGs from wildtype and tamoxifen-treated co-activator-expressing mice were isolated, challenged with LPS, and the transcriptome was profiled as before (**Fig. 3A**). PCA clustering again resolved LPS as the major axis of variation, with the co-activator and wildtype LPS samples partially separated (**Fig. 4D**). As with the primary human material, low RNA input limited the reliability of individual extreme fold-change estimates, and interpretation was therefore anchored on pathway-level enrichment (**Supplementary Fig. 6A-C**). Unsupervised Reactome pathway analysis revealed that the most significantly induced pathways in co-activator-expressing microglia were mitochondrial and oxidative, including Complex I Biogenesis, Respiratory Electron Transport, and Aerobic Respiration, while suppressed pathways included neuronal-signaling and cell-cell-communication modules (**Fig. 4E**). The targeted Hallmark analysis showed negative enrichment across all seven inflammatory pathways, with the strongest, statistically significant suppression in the interferon alpha response and TNFα signaling via NF-κB, and a negative trend in the inflammatory-response pathway (**Fig. 4F**).

qPCR confirmed a significant, gene-dosage-dependent increase in *Pgc1α* transcript (up to approximately tenfold over wild type) and a significant corresponding gene-dosage-dependent reduction in LPS-induced *Tnfα*, decreasing from wildtype to heterozygous to homozygous synthetic transcriptional co-activator microglia (**Fig. 4G**). Consistent with the induced mitochondrial biogenesis program, qualitative Airyscan super-resolution imaging of MitoTracker-labeled primary retinal microglia revealed a pronounced expansion of the mitochondrial network in *Activ_PGC1α*-expressing cells, which displayed a denser, more elaborate mitochondrial architecture than the sparse, punctate mitochondria of wildtype controls, providing organelle-level confirmation that the transcriptional oxidative phosphorylation signature translates into increased mitochondrial mass (**Supplementary Fig. 7A**). The convergence of this orthogonal genetic gain-of-function on PGC1α with the ZLN005 pharmacology on the same anti-inflammatory signature argues that the shared effect is attributable to PGC1α itself rather than to an off-target activity of the small molecule. In addition, whereas acute ZLN005 agonism mainly engaged the anti-inflammatory arm, our transcript data suggests that sustained expression of *Activ_PGC1α* engaged both the transcriptional and the oxidative arms simultaneously.

### Microglia-specific *Activ_PGC1α* expression reduces retinal degeneration in a humanized ***RHO^P347L/+^* mouse model of autosomal dominant RP**

To test therapeutic potential in vivo, we crossed the co-activator line to our humanized *RHO^P347L^* model of adRP ^25^, generating *Tmem119^CreERT^*^2^; *R26^LSL-Activ_PGC1α/+^*; hu*RHO^P347L^/*mu*Rho^WT^*animals (**Fig. 5A**). Because the *R26* and *Rho* loci are syntenic and the hu*RHO^P347L^* allele must be maintained heterozygous to allow a meaningful window of analysis, all experimental animals were heterozygous for both alleles and the reported effect sizes therefore likely reflect a lower bound. Tamoxifen was administered at postnatal day 21 to induce microglia-restricted *Activ_PGC1α* expression, with uninjected littermates of the same genotype as controls. Because P21 induction coincides with the earliest signs of degeneration, this design models a clinically more relevant post-onset intervention rather than a pre-symptomatic regimen.

**Fig. 5.**
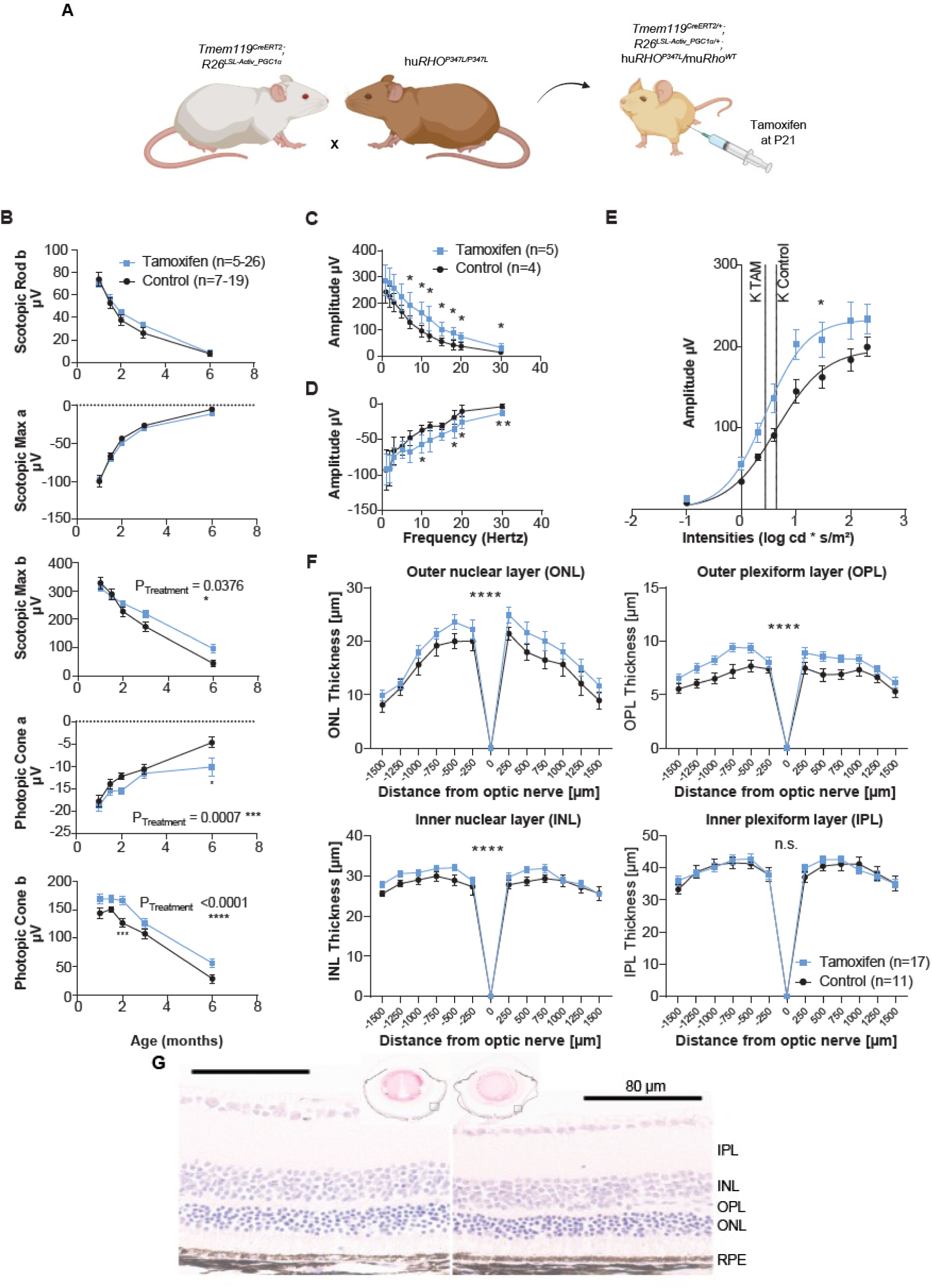
Microglia-restricted *Activ_PGC1α* expression retarded retinal degeneration in a humanized *RHO^P347L/+^*mouse model of autosomal dominant retinitis pigmentosa. (**A**) Breeding and induction scheme: *Tmem119^CreERT2^*; *R26^LSL-Activ_PGC1α/+^*; hu*RHO^P347L/+^* animals received tamoxifen at postnatal day 21 to drive microglia-restricted *Activ_PGC1α* expression; uninjected littermates served as controls. (**B**) Longitudinal electroretinography (ERG) from 1 to 6 months. Photopic cone responses show the most consistent preservation (two-way ANOVA a-wave *P*Treatment = 0.0007; b-wave *P*Treatment < 0.0001) and mixed rod-cone b-wave amplitudes are partially preserved (*P*Treatment = 0.0376), whereas pure-rod and mixed a-wave amplitudes are not. (**C, D**) Dark-adapted flicker ERG at 2 months: Activ_PGC1α animals show higher b-wave amplitudes across the cone-relevant flicker range (5-30 Hz) and higher a-wave amplitudes at high frequencies. (**E**) Naka–Rushton fit of the light-adapted (photopic) b-wave intensity– response relationship (extra sum-of-squares *F*(3,66) = 10.71, *P* < 0.0001); the tamoxifen group shows higher maximal response (*R*max 233.3 vs 196.9 µV) and a leftward-shifted semi-saturation constant (*K* 2.739 vs 4.369). (**F**) Retinal-layer thickness versus eccentricity from H&E sections; the outer nuclear, outer plexiform and inner nuclear layers are preserved (all *P*Treatment < 0.0001, two-way ANOVA), whereas the inner plexiform layer is unchanged. (**G**) Representative H&E retinal cross-sections (tamoxifen - left versus control - right). Scale bar, 80 µm. Data are mean ± SEM; replicate numbers are indicated.

Longitudinal electroretinography from one to six months revealed progressive, durable preservation of cone-driven function (**Fig. 5B**). Photopic cone responses showed the most consistent rescue (two-way ANOVA a-wave pTreatment = 0.0007; b-wave pTreatment < 0.0001), indicating durable protection of the cone pathway, followed by mixed rod-cone b-wave amplitudes (pTreatment = 0.0376). By contrast, pure rod and mixed rod-cone a-wave amplitudes were not preserved. This dissociation is consistent with microglia-restricted *Activ_PGC1α* expression not being designed to prevent the cell-autonomous rod dysfunction caused directly by the hu*RHO^P347L^*trafficking defect, but to dampen the secondary, inflammation-associated damage that follows rod death. Additional dark-adapted flicker ERGs and light-adapted serial-intensity ERGs were recorded at two months in a separate litter **(Fig. 5C–E)**. *Activ_PGC1α*-expressing animals showed significantly higher b-wave amplitudes across the cone-relevant flicker range (for 7-30 Hz) and higher a-wave amplitudes at the high-frequency end, consistent with preserved responsiveness across the tested flicker frequencies. Naka-Rushton fitting of the photopic b-wave intensity-response relationship revealed significantly different curves between groups (extra sum-of-squares F(3,66) = 10.71, p < 0.0001), with the tamoxifen group showing higher maximal response (Rmax 233.3 µV, 95% CI 211.7-263.8) and a leftward-shifted semi-saturation constant (K 2.739, 1.924-4.261) relative to controls (Rmax 196.9 µV, 179.3-224.2; K 4.369, 3.117-7.241), consistent with enhanced photopic responsiveness (**Fig. 5E**).

Structural analyses paralleled the functional rescue (**Fig. 5F, G**). H&E-stained retinal cross-sections revealed preserved thickness across eccentricity in *Activ_PGC1α*-expressing mice for the outer nuclear layer, the outer plexiform layer, and the inner nuclear layer (all pTreatment < 0.0001, two-way ANOVA), whereas the inner plexiform layer was unchanged between groups (**Fig. 5F**). This indicates that the site of Activ_PGC1α-mediated protection is spatially concentrated to the more outer retina where inflammation-associated microglial migration and subretinal accumulation occur, and where rod photoreceptor loss originates. Analysis of RPE flatmounts stained for the tight-junction marker ZO-1 revealed more homogenous, significantly larger, and more uniform RPE cell area (p < 0.001) and higher RPE circularity (p < 0.001) in tamoxifen-injected animals relative to the heterogeneous, patchy morphology of uninjected disease controls, indicating preservation of the regular hexagonal RPE monolayer that deteriorates during photoreceptor-degeneration-associated RPE stress ^28^ (**Supplementary Fig. 8A-C**). Comparative assessment of PNA-labeled cone photoreceptors in retinal flatmounts of the same litter revealed a near-significant trend toward greater cone-occupied area in tamoxifen-injected animals (p = 0.072), directionally consistent with the functional preservation observed by electroretinography (**Supplementary Fig. 8D, E**).

Quantitative morphometry of IBA1-stained subretinal cells revealed that tamoxifen-injected animals showed a significant reduction in IBA1^+^ cell solidity (p < 0.01), reflecting a shift from compact, amoeboid morphology toward ramified processes, and a significant increase in the number of processes per cell (p < 0.01), indicating more homeostatic microglial phenotypes (**Supplementary Fig. 9A, B**). The number of IBA1^+^ cells in the subretinal space, normalized to imaged area, did not differ between groups, indicating that the rescue is not attributable to reduced microglial recruitment to the outer retina. Because Müller glia and microglia participate in a reciprocal reactive-gliosis loop, we assessed Müller reactivity by quantitative GFAP immunostaining. GFAP-positive area was reduced in co-activator-expressing animals relative to controls (Unpaired student’s t-test, p-values ranging from 0.0158 to 0.0262, for 10-25 µm process-length cut-off), with mean process length differing significantly across all cut-offs examined (p < 0.05 at all cut-offs), with mean process length differing significantly across all cut-offs examined (p < 0.05 at all cut-offs), whereas total process length showed a consistent reduction that reached significance at cut-offs ≥19.5 µm (p = 0.034–0.0498). This length-dependent weakening of Müller gliosis suggests attenuation of the reactive microglia-Müller loop in Activ_PGC1α retinae (**Supplementary Fig. 9C, D**).

In summary, microglia-restricted *Activ_PGC1α* expression is sufficient to attenuate multiple hallmarks of RP pathology in vivo, including cone functional decline, outer-retinal structural degeneration, RPE disruption, and microglial and Müller-glia activation. This establishes a non-cell-autonomous neuroprotective role for microglial metabolic reprogramming in inherited retinal degeneration.

## Discussion

Retinitis pigmentosa is genetically heterogeneous, yet its downstream pathophysiology converges on microglia-driven neuroinflammation that operates largely independent of the initiating mutation. Building on human evidence that inflammatory signaling is pharmacologically engageable in RP ^7,8^, we asked whether the cellular source of RP-linked inflammation, the retinal microglial, can be stabilized to reduce disease severity.

Recent work reported that intravitreal ZLN005 rescues photoreceptor function in the rd1 mouse model and attributed the benefit to photoreceptor-autonomous mitochondrial biogenesis via the PGC1α/NRF-1/TFAM axis ^18^. Because resident microglia are the mediators of the inflammatory milieu that drives secondary photoreceptor loss ^29–31^, we reasoned that they are a plausible cellular target. We therefore used human iPSC-derived microglia as a scalable, tractable system for probing microglia-intrinsic responses to inflammatory and pharmacological challenges ^32^.

Transcriptionally, the dominant footprint in ZLN005-treated iMGs after 6 h of LPS is *IRF/STAT/ISG* collapse, with NF-κB-driven cytokines reduced at the protein level after 24 h. Type-I and -II interferon and STAT1-linked signaling have independently been implicated as photoreceptor-toxic microglial programs in retinal degeneration ^12,13^. Cross-platform confirmation in primary donor human retinal microglia showed the same directionality, positioning resident retinal microglia as plausible effectors of PGC1α-directed protection and as target for intervention.

PGC1α-mediated restraint of NF-κB and IFN signaling likely operates through multiple complementary routes. Direct p65 interaction was observed in cardiac cells ^33^; enhanced mtDNA quality control that dampens cytosolic cGAS activation ^34^, consistent with the *SIRT1* transcript increase and the *cGAS/STAT1/2/ISG* induction in our LPS contrast and collapse under ZLN005; and co-activation of PPARα and ERRα, both of which transrepress NF-κB and TLR4 signaling ^35,36^, consistent with *PPARα* and *ERRα* induction in our dataset.

Against this consistent anti-inflammatory backdrop, an unexpected feature emerged. The oxidative arm classically coupled to PGC1α agonism failed to engage under concurrent TLR4 signaling, even though it was intact under homeostatic conditions. This override is unlikely to reflect insufficient exposure time, as ZLN005 elevates respiration in retinal pigment epithelial cells over a comparable exposure ^37^, placing our 48 h pre-conditioning window within the expected range. Under LPS co-stimulation, mitochondrial ATP production fell just below the vehicle control rather than trending toward a delayed rise, and lactate remained elevated, a signature of a sustained TLR4 signaling-associated metabolic phenotype. This barrier was also visible directly in our carbon flux assay. Rather than a "broken TCA cycle" ^38^, ¹sC incorporation into citrate, glutamate, and malate increased under LPS, so the elevated lactate to citrate ratio reflects glycolytic overflow past an active cycle ^39^, with the oxidative arm likely outrun rather than switched off. Carbon tracing further revealed engagement of pyruvate carboxylase-mediated anaplerosis under LPS (M+3/M+2 aspartate ratio 0.28 → 0.43), with aspartate pool contraction relative to other TCA metabolites despite increased anaplerotic replenishment, a pattern consistent with cataplerotic demand for aspartate (e.g., argininosuccinate synthesis for nitric oxide production) rather than respiratory constraint ^40^. This pattern matches human iPSC-derived microglia, in which inflammatory activation preserves mitochondrial respiration and raises TCA cycle enzyme abundance, unlike rodent microglia ^41^. The anti-inflammatory action therefore might instead operate through a PGC1α-mediated restraint of NF-κB and STAT signaling ^17,42^, that is at least partially uncoupled from acute oxidative reprogramming, consistent with recent evidence that pro-inflammatory macrophage activation does not require inhibition of oxidative phosphorylation ^43^.

Activ_PGC1α was designed to mitigate this specific mechanistic dissociation via enhanced output-per-molecule. Because inflammation post-translationally suppresses endogenous PGC1α activity while diverting metabolism away from oxidative output ^16,22^, upstream pharmacological stimulation cannot escape the inflammatory ceiling. Raising per-molecule transcriptional co-activator output through a fused transactivation domain acts downstream of that suppression node. In line, sustained genetic activation in primary retinal microglia engaged both the anti-inflammatory and oxidative arms under LPS with coordinate induction of oxidative-phosphorylation programs and negative enrichment of all targeted LPS-response pathways. Tethering the TAD stack to a native co-activator confines hyperactivation to its partner-receptor repertoire, avoiding the sgRNA-independent off-target activation and co-factor squelching documented for dCas-based TAD delivery ^23,44^. The gene-dosage-dependent *Tnfα* attenuation across wildtype, heterozygous, and homozygous microglia demonstrates that the achievable engagement is therapeutically consequential and thus TUNE represents a viable strategy for targeted pathway engagement. Our efforts are focused on validating these results in non-cancerous human microglia; however, this is currently hampered by difficulties in introducing large transgenes into primary or iPSC-derived microglia.

Microglia-restricted *Activ_PGC1α* expression, induced at P21 in our hu*RHO^P347L^* model, conferred durable functional and structural preservation consistent with reducing secondary degeneration rather than correcting the primary rod defect. The model carries the full human *RHO P347L* sequence at the endogenous mouse locus and can support combination studies with gene-directed interventions ^25,45,46^. Its heterozygous state recapitulates autosomal-dominant inheritance and creates a chronic window for longitudinal post-onset testing. This design therefore addresses a distinct therapeutic component: limiting mutation-independent inflammatory damage after rod dysfunction begins.

The ONL rescue that we observed indicates that rod cell bodies were also retained despite the absent a-wave rescue, as cone photoreceptors constitute approximately 2 to 3% of total photoreceptors in the mouse retina ^47^ and therefore cannot attribute to the observed ONL rescue alone. We attribute this retention to relief of the inflammatory microglial pressure that would otherwise clear the surviving cells. Microglial IL-1β amplifies photoreceptor apoptosis directly and through Müller-cell chemokine relay ^29,30^, activated microglia phagocytose stressed but viable photoreceptors displaying phosphatidylserine "eat-me" signals ^29^, and ONL-infiltrating microglia inflict complement-associated damage ^31^. The observed protection occurred without measurably reducing the density of subretinal IBA1+ cells, arguing that the therapeutic action is a change of state imposed on relocated microglia rather than suppression of infiltration itself. Consistent with a state-level effect propagating through the reactive-gliosis network, quantitative GFAP analysis showed reduced Müller-glial reactivity in co-activator-expressing retinae. Because microglial IL-1β drives Müller-cell chemokine relay ^29,30^, attenuation of Müller gliosis provides independent anatomical evidence that microglial reprogramming weakens the reciprocal microglia-Müller reactive loop rather than acting on microglia alone.

Our findings extend prior work on microglial PGC1α. Total monocyte-specific (Cx3cr1-Cre) elevation of native PGC1α attenuates cytokine output and NLRP3 activation in acute stroke ^17^. We extend this in three ways: validation across human iPSC-derived, primary donor retinal, and primary mouse retinal microglia with a microglia-specific driver ^48^, identification of interferon, NF-κB, IL6/JAK/STAT3, and complement signaling as the dominant suppressed programs rather than NLRP3, and durable rescue over six months in a chronic genetic degeneration model. Finally, in the PPRE reporter system, our TUNE’d PGC1α system achieves 2.3-to 2.5-fold higher output than native PGC1α, validating that transactivation-domain engineering can raise transcriptional co-activator output beyond endogenous levels.

Several limitations deserve emphasis. First, *R26-Rho* synteny forced heterozygous Activ_PGC1α dosing in vivo, likely underestimating the achievable benefit, a limitation supported by the gene-dosage-dependent *Tnfα* decrease from wildtype to heterozygous to homozygous microglia. Second, low post-mortem retinal microglia yield, approximately 0.3% and 0.1% of human and murine retinal cells, respectively ^9,49^, constrained donor-paired transcriptomics and prevented respirometric confirmation in primary rMGs. Third, *Tmem119^CreERT2^* is active in CNS microglia broadly, so a brain-retina axis contribution cannot be excluded. Fourth, our flux inferences derive from a ¹sC-glucose tracer and thus capture only the glucose-derived fraction of each pool. Because TCA intermediates in activated microglia are predominantly filled by non-glucose carbon, glutamine and fatty-acid contributions to the observed ratio shifts were not directly resolved and require further investigation. Finally, LPS engages TLR4 signaling whereas RP is driven by sterile DAMPs, such as oxidized outer-segment lipids, HMGB1, mtDNA-cGAS, complement ^50^. However, these triggers converge on the same NF-κB, IRF3/STAT1, and type-I IFN nodes PGC1α restrains ^51^, with the hu*RHO^P347L^*model supplying relevant in vivo validation.

In summary, by combining pharmacological and genetic perturbations across human iPSC-derived, primary human donor, and humanized mouse microglia, we show that PGC1α limits inflammatory activation in human microglia without restoring their oxidative capacity, and that microglia-restricted expression of an engineered hyperactive PGC1α co-activator, is sufficient to confer durable, non-cell-autonomous neuroprotection in a humanized model of autosomal-dominant retinitis pigmentosa. This identifies microglial immunometabolism as a tractable, mutation-agnostic therapeutic axis that complements rather than competes with gene-replacement and gene-editing strategies, and establishes synthetic transcriptional co-activators as a payload class for cell-type-restricted neuroprotection.

## Methods

### Preclinical models

All mice were housed in the Columbia University Pathogen-Free Eye Institute Animal Facility and maintained with a 12 h light/12 h dark cycle with access to food and water ad libitum. At indicated timepoints, mice were euthanized by CO_2_ before cervical dislocation.

hu*RHO^P347L^* mice were generated in-house by HDR-mediated humanization of the endogenous mouse *Rho* locus using two SpCas9 gRNAs and a single-stranded oligonucleotide donor encoding the full human *RHO* sequence at the endogenous mouse locus with the disease-causing P347L mutation (c.1040C>T). Founder animals were backcrossed three times to C57BL/6J. Full generation, backcross, genotyping, and characterization details are provided in a companion manuscript ^25^. All mice were verified negative for common *rd1/rd2/rd8/rd10* alleles. *R26^LSL-Activ_PGC1α^* mice carried a conditional expression cassette targeted to the *Rosa26* locus. The knock-in cassette comprised, in 5′-to-3′ orientation: the CAG promoter; a loxP-flanked transcriptional STOP cassette containing the PGK-neomycin-resistance gene followed by three tandem SV40 polyadenylation signals; the *Activ_PGC1α* transgene; and a bovine growth-hormone polyadenylation signal. The triple-polyadenylation STOP cassette prevents transcriptional read-through in the absence of Cre. Activ_PGC1α encoded an N-terminal multivalent p65-HSF1 ^26^ transactivation module directly fused to murine *PGC1α* (*Ppargc1a*; RefSeq NM_008904.3). The complete nucleotide and predicted amino-acid sequences, including domain order, orientation, and the *Rosa26*, cassette recombination junction, together with targeted-allele validation data are provided in Supplementary Table 2 and Supplementary Fig. 10. *R26^LSL-Activ_PGC1α^*, *Tmem119^CreERT2^* mice were derived by crossing to C57BL/6-*Tmem119^em1(cre/ERT2)Gfng/J^* (JAX #031820) ^48^. Experimental animals (*R26^LSL-Activ_PGC1α/+^*, *Tmem119^CreERT2/+^*, hu*RHO^P347L^*/mu*Rho^WT^*) were generated by further crossing with hu*RHO^P347L^*/hu*RHO^P347L^*mice, with littermates used for comparisons. Tamoxifen (100 mg/ml in ethanol; Sigma-Aldrich) was diluted to 10 mg/ml in sunflower seed oil and injected i.p. at P21 for three consecutive days (0.1 mg/g), followed by a 10-day interval before experiments.

### Primary mouse retinal microglia culture

Eyes were enucleated from euthanized mice of both sexes, cleaned of extraocular muscles, and gently perforated. Both eyes from each animal were placed onto a 25 cm² tissue culture plate for 1 min before addition of 3 mL pre-warmed DMEM supplemented with 10% FBS, 1% Antibiotic-Antimycotic (Gibco), and 25 ng/mL M-CSF1 (PeproTech) ("primary culture medium"). Cultures were maintained until confluent with medium changes every 7 days.

### Primary human donor retinal microglia culture

Human donor ocular tissues were obtained from The Eye-Bank for Sight Restoration, Inc. (New York, NY) as fully anonymized, de-identified specimens in compliance with federal privacy regulations. Eyecups were obtained from deceased donors within 8 h post mortem. Whole globes were transported on ice in DMEM. Under sterile conditions, anterior segments were removed and posterior eyecups were prepared by four radial incisions from the peripheral sclera toward the optic nerve head. The vitreous was removed, and the neural retina was gently separated from the RPE by blunt dissection. Retinal tissue was sectioned and placed onto 25 cm² plates for 1 min before addition of 3 mL primary culture medium. Cultures were maintained until confluent with medium changes every 7 days. Donor details are provided in Supplementary Table 1.

### Induced pluripotent stem cell (iPSC)-derived microglia culture

Human iPSCs were differentiated into microglia (iMG) using the STEMdiff™ Hematopoietic Kit (Catalog #05310) and STEMdiff™ Microglia Differentiation and Maturation Kits (Catalog #100-0019 and #100-0020) per manufacturer’s instructions. huPSCs were directed toward hematopoietic progenitors over 12 days, after which 150,000-200,000 floating progenitors were plated onto Matrigel-coated plates in differentiation medium for 23 days, followed by maturation medium. A subset of cells was routinely used to verify cellular purity via CD11b^+^ flow cytometry, confirming 90%-96% CD11b^+^ cell purity. Experiments were performed after 2 days in maturation medium.

### HEK cell culture

HEK293T cells were cultured in DMEM (Gibco) supplemented with 10% FBS (Gibco) at 37 °C with 5% CO2. Cell lines tested negative for mycoplasma.

### Analysis of PGC1α-driven suppression of LPS responses *in vitro*

Cultures were divided into four groups with at least three biological replicates: [1] vehicle control, [2] 1 µg/mL LPS (eBioscience™) for 6 h (bulkRNAseq) or 24 h (ELISA and metabolomics), [3] 20 µM ZLN005 (Sigma-Aldrich) for 48 h, and [4] ZLN005 for 48 h with LPS for the final 6 h (bulkRNAseq) or 24 h (ELISA and metabolomics). Mutant mouse retinal cultures were treated accordingly without ZLN005. All conditions received equivalent total medium volume per well. Cells were lifted using cell scrapers; primary retinal cells were purified via FACS, while iMG were directly collected into TRIzol (Invitrogen) and stored at −80 °C.

### FACS isolation of retinal microglia

Lifted cells were filtered through 100 µm cell strainers (Corning), pelleted at 300 × g for 5 min at 4°C, and resuspended in ice-cold HBSS + 2% FBS. After Fc-block (TruStain FcX™, Biolegend) for 10 min, cells were incubated with anti-CD11b antibody (FisherScientific, MABF512MI) for 30 min, washed, and resuspended in HBSS with SYTOX blue viability dye. Single, viable CD11b^+^ cells were sorted into TRIzol (Invitrogen) on a Sony MA900 cell sorter. For rMG analysis, 8-10 primary rMG cultures each derived from separate mice and treated equally were pooled for a single datapoint.

### RNA isolation and quantification

RNA was isolated using the Direct-zol RNA MiniPrep Plus kit (Zymo Research) with on-column DNase digestion. cDNA was generated with LunaScript RT Master Mix (NEB). qPCR was performed using Fast SYBR Green Master Mix (Sigma) on a QuantStudio 5 (ThermoFisher Scientific). *Rps18* or *ACTB* served as housekeeping genes for mouse or human samples, respectively, selected by lowest coefficient of variation across all samples. Primer sequences are listed in Supplementary Table 3. One-way ANOVA with Tukey’s test was used for statistical evaluation (GraphPad Prism V8.4).

For bulk-RNAseq, RNA concentration and integrity were verified by Qubit and Tapestation 4200. Low-input samples (<100 ng) were processed with SMART-Seq v4 Ultra Low Input RNA Kit (Takara) followed by Nextera XT (Illumina; 40M reads). Higher-input iPSC samples were processed with TruSeq Stranded mRNA Library Prep Kit (Illumina; 40M reads) with KAPA HiFi HotStart Ready Mix for the final PCR. Libraries were sequenced on the Element Aviti at the Columbia Sulzberger Genome Center. Reads were quantified at the transcript level using kallisto.

### Bulk-RNA sequencing

Transcript-level abundances were quantified using kallisto (v0.46.1) against Ensembl GRCm39 (mouse) or GRCh38 (human) reference transcriptomes. Gene-level counts were imported into R for downstream analysis. Differential expression was performed using DESeq2 (v1.46.0).

The iMG dataset comprised three conditions (Control, Control_LPS, ZLN005_LPS; n = 3 per group; total n = 9), modeled with ∼condition, contrasting ZLN005_LPS vs Control_LPS. The human donor rMG cohort comprised four donors profiled under LPS and LPS+ZLN005; one donor was excluded as a PC1 outlier with reduced library complexity, yielding n = 6 (3 donors × 2 conditions), modeled with ∼donor + condition, contrasting LPS+ZLN005 vs LPS. The mouse rMG dataset comprised four conditions (Wildtype_Control n = 3, Activ_PGC1α_Control n = 3, Wildtype_LPS n = 3, Activ_PGC1α_LPS n = 4), modeled as ∼group, contrasting Activ_PGC1α_LPS vs Wildtype_LPS.

Genes with fewer than 10 summed raw counts were excluded. Library-size normalization used DESeq2’s median-of-ratios method. Dispersion estimates used the default parametric fit with empirical Bayes shrinkage. Hypothesis testing used the Wald test with Benjamini-Hochberg FDR correction. Genes were called differentially expressed at FDR < 0.05 and |log2FC| ≥ 1. log2FC shrinkage via apeglm (v1.28.0) was applied post hoc for ranking and MA plots. Volcano plots display unshrunken log2FC and −log10(FDR), the top 15 DEGs were labeled.

For quality assessment, rlog-transformed counts were used for PCA and heatmaps of the 500 most variable genes (1 − Pearson correlation, Ward.D2 linkage, z-scored per gene). Heatmaps used ComplexHeatmap (v2.22.0), volcano plots used ggplot2 (v4.0.3).

Pathway analysis used pre-ranked GSEA with fgseaMultilevel (fgsea v1.32.2; minSize = 15, maxSize = 500, eps = 0, seed = 42). Hallmark (H) and Reactome (C2:CP:REACTOME) gene sets were retrieved from MSigDB v2026.1 via msigdbr (v26.1.0), using mouse or human collections as appropriate. Pathways with BH-adjusted p < 0.05 were considered significant. To quantify LPS-responsive inflammatory reprogramming, NES values for seven predefined Hallmark gene sets (TNFA_SIGNALING_VIA_NFKB, INFLAMMATORY_RESPONSE, IL6_JAK_STAT3_SIGNALING, INTERFERON_ALPHA_RESPONSE, INTERFERON_GAMMA_RESPONSE, COMPLEMENT, APOPTOSIS) were extracted for cross-dataset comparison.

For human iMGs, a pre-specified 25-gene panel spanning six functional themes (mtDNA-QC, cGAS-STING, JAK-STAT, ISGs, IRF chemokines, nuclear receptors) was visualized per condition as VST-based box-and-whisker plots with overlaid sample points using ggplot2 (v4.0.3) with the ggprism (v1.0.7) theme. Bracketed significance stars for the Control_LPS vs Control and ZLN005_LPS vs Control_LPS contrasts report the same DESeq2 Wald-test Benjamini-Hochberg-adjusted p-values used throughout (**** ≤ 0.0001, *** ≤ 0.001, ** ≤ 0.01, * ≤ 0.05).

### L-Lactate quantification from culture supernatants

Culture supernatants were collected, centrifuged at 300 × g for 5 min, and stored at −80 °C. L-lactate was measured using the L-Lactate Assay Kit (Eton Bioscience; Fisher Scientific #NC0014474). Supernatants were incubated with L-Lactate Assay Solution for 30 min at 37 °C, stopped with 0.5 M acetic acid, and absorbance read at 490 nm on a SpectraMax M5. Concentrations were interpolated from a standard curve run in parallel. The same supernatants were used for cytokine ELISAs.

### Protein isolation and quantification

Human Luminex® Discovery Assays for IFN-gamma, IL1-beta, IL-6, IL-10, and TNF-alpha were performed on culture supernatants per manufacturer’s instructions and measured on a Luminex 200 system. One-way ANOVA with Tukey’s test was used for statistical evaluation (GraphPad Prism 8), values below the lower limit of detection were set to 0.

### Seahorse XF Real-Time ATP Rate Assay

OCR and ECAR were measured on a Seahorse XFe96 Analyzer (Agilent) using the XF Real-Time ATP Rate Assay Kit (#103592-100). iMG were seeded onto CellTak-coated XFe96 plates at 150,000 cells/well two days before assay and treated as before (Control n = 3, ZLN005 n = 6, LPS n = 3, ZLN005+LPS n = 6). On the day of measurement, medium was replaced with XF DMEM (pH 7.4) supplemented with 10 mM glucose, 1 mM pyruvate, and 2 mM glutamine, and equilibrated for 1 h at 37 °C without CO2. After three baseline measurements, oligomycin (1.5 µM) was injected, followed by rotenone/antimycin A (0.5 µM each). mitoATP and glycoATP production rates were calculated from OCR and ECAR following the XF Real-Time ATP Rate Assay equations: mitoATP = (mean basal OCR − mean oligomycin-insensitive OCR) × 2.75; glycoATP = (PER − mitoPER) × 2.75, with PER = ECAR × 7.394 (buffer factor × chamber volume × Kvol) and mitoPER = basal OCR × 0.61; PER was derived from ECAR for all wells. Basal OCR and ECAR were means of pre-oligomycin measurements. Statistical comparisons used one-way ANOVA with Tukey’s HSD.

### 13C isotopologue tracing

Mature iMG were seeded at 150,000 cells/well (n = 5-6 per condition) in STEMdiff™ Microglia Maturation medium with 20 mM 12C-glucose and 100 µM 12C fatty acid-BSA for 48 h; 20 µM ZLN005 was added for 48 h in respective groups and 1 µg/ml LPS for the final 24 h. Medium was then replaced for 6 h with tracer substrates: STEMdiff™ Microglia Maturation medium with 20 mM 13C-glucose with 100 µM 12C fatty acid-BSA. LPS and ZLN005 were maintained throughout. Supernatants were collected and cells scraped into 250 µl ice-cold 80% methanol with 10 µM methylsuccinate as internal standard. Extracts were analyzed by stable isotope-resolved metabolomics coupled to GC-MS ^45,52^. Dried extracts were derivatized with methoxyamine-HCl in pyridine and TBDMS. Isotopologue abundances were quantified using MSD ChemStation, and corrected for natural isotopic abundance using IsoCor.

Total-metabolite pool ratios (lactate/citrate, aspartate/glutamate, lactate/pyruvate) were computed per biological replicate from natural-abundance-corrected total pools (labeled plus unlabeled) and are reported as mean ± s.d. (n = 6 for Control, LPS and LPS+ZLN005; n = 5 for ZLN005 alone). Each ratio was compared across conditions by one-way ANOVA with Tukey’s multiple comparisons test using GraphPad Prism 8.4.

Carbon partitioning was resolved at the level of individual metabolite nodes on a custom schematic of glycolysis and the first turn of the TCA cycle. Mass isotopologue distributions were acquired by GC-MS and corrected for natural isotope abundance using IsoCor. From the corrected distributions, two parameters were computed per node. The first was fractional ¹sC enrichment, defined as labeled signal divided by the total (labeled plus unlabeled) pool. The second was total metabolite pool size.

For each node, the diagnostic isotopologue was the species expected from [U-¹sC]glucose via glycolysis and one TCA turn. This was the M3 species for glyceraldehyde-3-phosphate/DHAP, 3-phosphoglycerate, phosphoenolpyruvate, pyruvate and lactate, and the M2 species for citrate, glutamate, succinate, fumarate, malate and aspartate. For the symmetric C4 intermediates (succinate, fumarate, malate) and aspartate, M2 was retained as the diagnostic species with the understanding that fumarase-mediated symmetry scrambles label between the C1 to C2 and C3 to C4 positions, so enrichment values at these nodes reflect summed positional isotopologues rather than a single carbon origin.

Fractional enrichment (%, upper bar chart) and total pool size (lower bar chart) are displayed at each node as horizontal bars for the four conditions (mean ± s.d., n = 6 for Control, LPS and LPS+ZLN005, n = 5 for ZLN005 alone), with individual replicates overlaid. For each node, fractional enrichment and pool size were each compared between conditions by two-sample t-tests (all nodes, both metrics, and prespecified contrasts: LPS vs Control, ZLN005 vs Control, ZLN005 vs LPS+ZLN005m and LPS+ZLN005 vs LPS), and the resulting P values were corrected for multiple comparisons across all node-level tests using the Benjamini-Hochberg procedure; q denotes these false-discovery-rate-adjusted P values.

### Conditioned-medium transfer assay

Donor iMG were stimulated with LPS alone or LPS preceded by 24 h ZLN005 pre-treatment. After 24 h of LPS stimulation, donor cells were washed and re-exposed to LPS alone for 24 h, ensuring collected supernatants reflected donor pre-conditioning rather than stimulus carryover. Supernatants ("Sup LPS" and "Sup LPS+ZLN") were centrifuged at 300 × g for 5 min and applied undiluted to naive recipient iMG from an independent differentiation for 6 h. Recipient iMG were lysed and RNA analyzed as above.

### Assessment of PGC1α variants

Co-activator strength was assessed by quantifying secreted NanoLuc luciferase (secNluc) from a PPARγ-responsive PPRE reporter (PPRE-pNL1.3; Addgene #84394; ^27^). HEK293T cells were transfected at 60-80% confluency with 0.6 µl Lipofectamine 2000 and 300 ng reporter and/or 300 ng PGC1α variant-P2A-eGFP plasmid. Supernatant was collected every 24 h up to 96 h, signal from P2A-eGFP expression served as normalization control. Luminescence was measured using Nano-Glo® Luciferase Assay (Promega) on a SpectraMax M5 with SoftMax Pro 5. One-way ANOVA with Tukey’s test was used for statistical evaluation (GraphPad Prism).

### Assessment of mitochondrial morphology and quantity in primary retinal microglia

Wildtype and mutant CD11b^+^ murine primary retinal cells were FACS-isolated into primary culture medium, seeded onto ibiTreat-coated µ-Slide 18-well chambers (ibidi), and maintained for 2 days. Mitochondria were labeled with 100 nM MitoTracker Green FM and Hoechst for 30 min at 37 °C. High-resolution live-cell imaging was performed on a Zeiss LSM900 confocal with Airyscan super-resolution. 3D object reconstruction was performed using Imaris v.11 (Oxford instruments) and signal threshold was automatically adjusted to ensure accurate object segmentation.

### Electroretinograms (ERGs)

ERGs were recorded from both eyes and averaged at 1, 1.5, 2, 3, and 6 months of age using previously described methods ^28,45^. Mice were dark-adapted for 12 h, anesthetized with ketamine/xylazine, and placed on heating pads at 37 °C. Pupils were dilated with 1% tropicamide; gonio hypromellose (2.5%) was applied for corneal protection. Recordings were acquired at 1000 Hz (20 ms pre-trigger, 200 ms post-trigger). Step 1: low-intensity white flash (0.001 cd·s/m²) at 0.2 Hz, 10 sweeps averaged. Step 2: moderate-intensity white flash (3 cd·s/m²) at 0.2 Hz, 5 sweeps averaged. Step 3: after 10 min light adaptation, high-intensity xenon flash (30 cd·s/m²), 20 sweeps averaged.

An additional full-field protocol was performed on 2-month-old littermates (tamoxifen n = 5, control n = 4): a dark-adapted flash intensity series (−6.0 to +1.0 log₁₀ cd·s/m²), a dark-adapted flicker series (0.5–30 Hz at +0.5 log₁₀ cd·s/m²), and a subsequent light-adapted flash intensity series (−1.0 to +2.3 log₁₀ cd·s/m²). The light-adapted b-wave intensity–response relationship was fitted with a Naka–Rushton function. Longitudinal ERG outcomes were analyzed by two-way repeated-measures mixed-effects analysis (Age, Treatment, Age×Treatment) with Sidak’s multiple comparisons test (control vs. tamoxifen at each age, corrected across the five ages).

### Assessment of cone outer segment, RPE and microglia density and morphology

Whole retinas and posterior eye cups were flat-mounted as described ^45^. Retinae were stained with fluorescein-conjugated peanut agglutinin (PNA; Invitrogen L21409, 1:350) for cone outer segments and anti-IBA1 (WAKO Fujifilm 019-19741, 1:1000) for microglia. RPE cell borders were stained with anti-zona occludens 1 (ZO-1, ThermoScientific #40-2200, 1:25). Z-stacks were acquired on a Nikon AXR resonant scanning confocal at 20× and stitched into maximum intensity projections.

Cone density was quantified as percentage of total retinal area occupied by PNA-positive signal in Fiji (ImageJ) using local Otsu thresholding (15-pixel neighborhood) and Gaussian blur (10-pixel radius). RPE morphology was assessed by manually tracing cell borders in 10 evenly dispersed regions (≥15 cells/region/flatmount) by two blinded analysts, quantifying area and circularity (4π·area/perimeter²). Subretinal IBA1^+^ cell density was quantified by automated thresholding and watershed segmentation (minimum area 30 µm²). Morphological characterization was performed in ≥10 randomly selected areas (≥10 cells/area) measuring soma size, process number, branch number, process length and thickness, and cell solidity (cell area/convex hull area).

### Assessment of GFAP-reactive gliosis

Frozen retinal cross-sections were stained for glial fibrillary acidic protein (GFAP; DAKO Z0334, 1:1000) to assess Müller-glial reactivity and imaged at 40× (0.2 µm/pixel) using a Nikon Eclipse TS2r epifluorescence microscope. GFAP-positive Müller cell processes were quantified in Fiji (ImageJ) by automated thresholding and particle analysis within defined regions of interest. Because gliosis readouts depend on the minimum process-length (Feret) cut-off applied during segmentation, we performed a sensitivity analysis across multiple Feret cut-offs (10-25 µm) and report the prespecified endpoint of GFAP-positive area (percentage of ROI) together with total and mean process length. Between-group comparisons (tamoxifen n = 7; non-injected controls n = 5) used unpaired Student’s t-test on animal-level values.

### Assessment of retinal layer thickness and immunohistochemistry of retinal cross-sections

Enucleated eyes were fixed in Z-Fix zinc formalin ^28^, embedded in paraffin, and sectioned at 5 µm through the optic nerve by a board-certified histologist. H&E staining and image acquisition were as described ^28^. Outer nuclear layer (ONL), outer plexiform layer (OPL), inner plexiform layer (IPL), and inner nuclear layer (INL) thickness were quantified at 250 µm intervals up to 1500 µm from the optic nerve on both sides. Statistical analysis used two-way ANOVA with Sidak’s multiple comparisons test.

## Data availability

The RNA-sequencing data generated in this study have been submitted to the NCBI Gene Expression Omnibus (GEO, accession GSE347438). All other data will be made available by the authors upon reasonable request.

## Acknowledgements

This research was funded in part through the NIH/NCI Cancer Center Support Grant P30CA013696 and used the Genomics and High Throughput Screening Shared Resource. These studies used the Confocal and Specialized Microscopy Shared Resource of the Herbert Irving Comprehensive Cancer Center at Columbia University, funded in part through the NIH/NCI Cancer Center Support Grant P30CA013696. The Columbia Ophthalmology Core Facility is supported by NIH Core grant 5P30EY019007

We thank the members of Jonas Children’s Vision Care (JCVC) and BME for sharing cell lines and ideas.

## Supplementary Information

**Supplementary Fig. 1.**
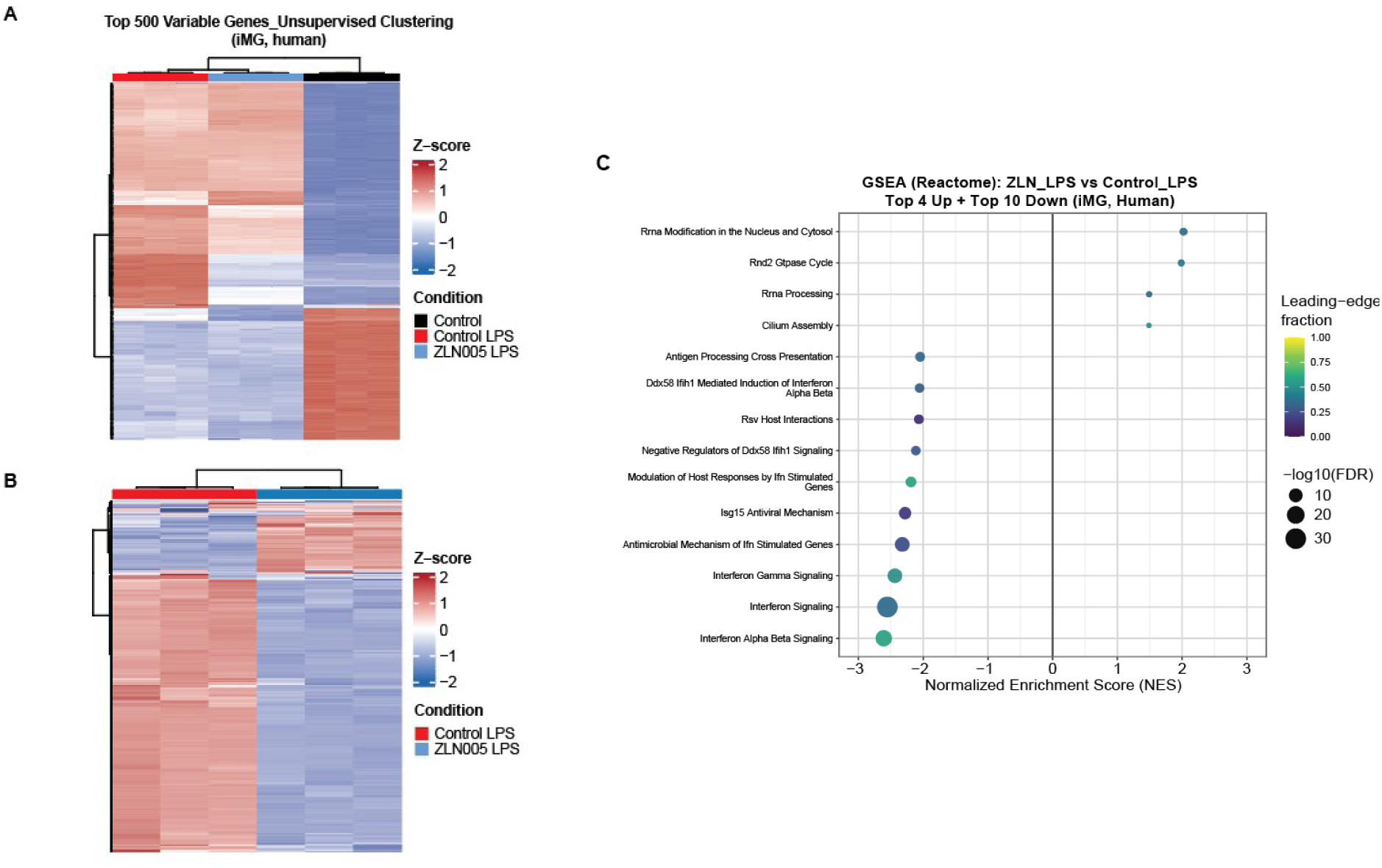
Transcriptional clustering and pathway enrichment in iPSC-derived microglia (iMG). (**A**, **B**) Unsupervised hierarchical clustering heatmaps of the top 500 most variable genes (row Z-scored): (**A**) across all three conditions (control, control+LPS, ZLN005+LPS) and (**B**) restricted to the control+LPS and ZLN005+LPS groups. (**C**) Reactome gene-set enrichment (GSEA) for the ZLN005+LPS versus control+LPS contrast (top significant 4 upregulated, top 10 downregulated pathways). The downregulated arm is dominated by interferon alpha/beta and gamma signaling and antiviral/ISG modules; positively enriched sets are non-inflammatory (rRNA processing, RND2 GTPase cycle, cilium assembly). Dot size, –log10(FDR); color, leading-edge fraction.

**Supplementary Fig. 2.**
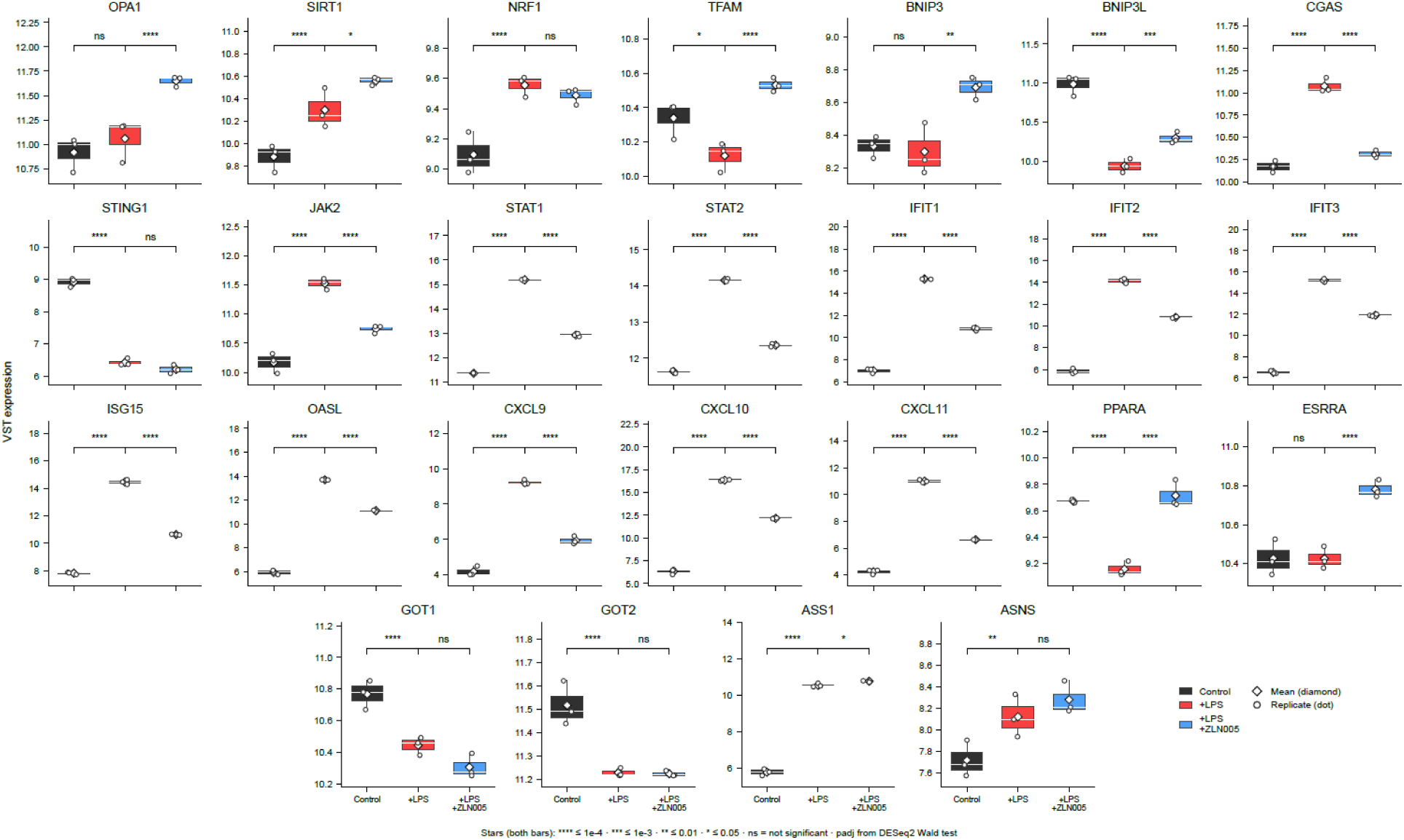
Gene-level expression of inflammatory, mitochondrial-biogenesis, and amino-acid–metabolizing transcripts in iMG. Boxplots of variance-stabilized (VST) expression for twenty-five transcripts across the three conditions (Control, +LPS, +LPS+ZLN005); boxes show the median and interquartile range, the white diamond marks the mean, and individual replicates are overlaid as points. Interferon-stimulated genes (*IFIT1, IFIT2, IFIT3, ISG15, OASL*), JAK–STAT components (J*AK2, STAT1, STAT2*) and inflammatory chemokines (*CXCL9, CXCL10, CXCL11*) are uniformly induced by LPS and significantly attenuated by ZLN005 co-treatment, whereas mitochondrial-biogenesis transcripts (*OPA1, SIRT1, NRF1, TFAM, BNIP3, BNIP3L*) show a mixed response. The cytosolic DNA sensor *CGAS*, but not STING1, is restored toward baseline by ZLN005. The PGC1α nuclear-receptor partners PPARA and *ESRRA* are induced by ZLN005 (*PPARA* is suppressed by LPS and restored toward baseline; *ESRRA* is unchanged by LPS and induced above it). Amino-acid–metabolizing enzymes are LPS-responsive: the aspartate aminotransferases *GOT1* and *GOT2* are downregulated, whereas *ASS1* and *ASNS* are upregulated, consistent with inflammatory aspartate depletion. Brackets denote the two prespecified contrasts (left, +LPS vs Control; right, +LPS+ZLN005 vs +LPS). Adjusted P values are from the DESeq2 Wald test (****P ≤ 1×10⁻⁴; ***P ≤ 1×10⁻s; **P ≤ 0.01; *P ≤ 0.05; ns, not significant).

**Supplementary Fig. 3.**
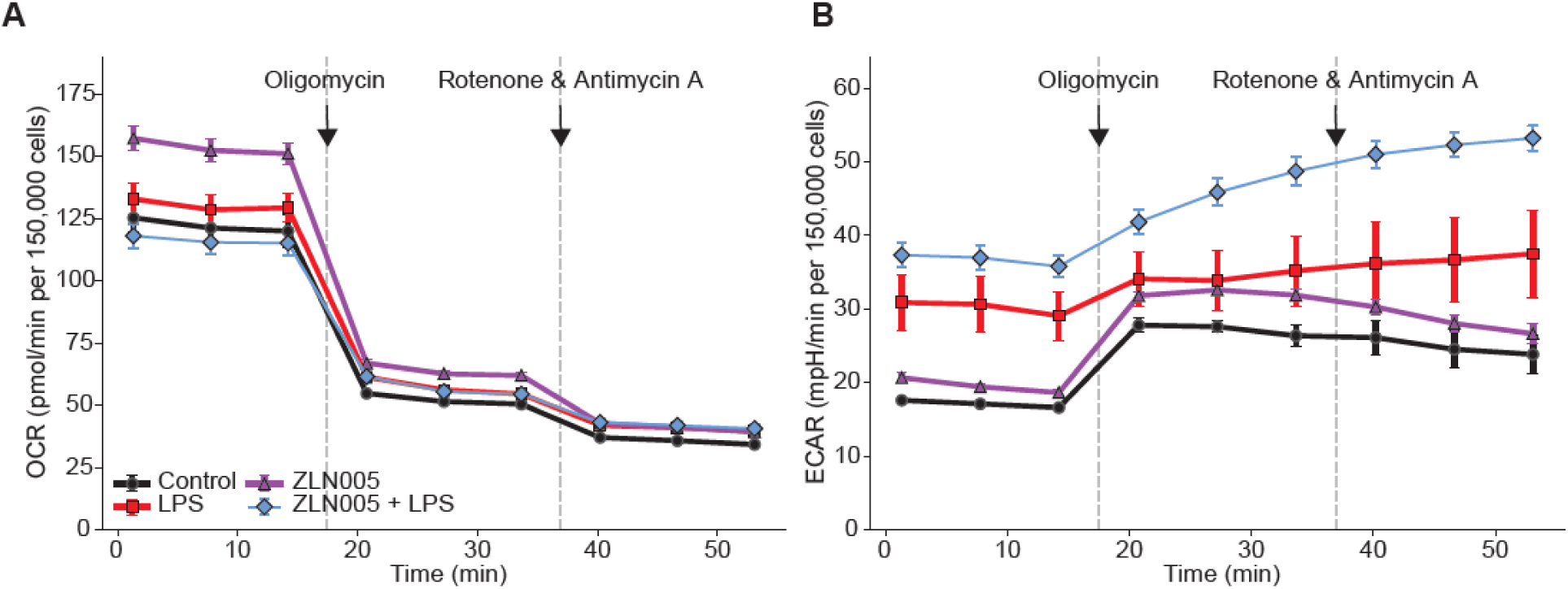
Real-time respirometry of iMG. (**A**) Oxygen consumption rate (OCR) and (**B**) extracellular acidification rate (ECAR) kinetic traces from the Seahorse ATP-rate assay across conditions, with sequential injection of oligomycin and rotenone + antimycin A (timing indicated). Cells were seeded at 150,000 cells/well and rates are expressed per 150,000 cells. n = 3 (Control, LPS) and n = 6 (ZLN005, ZLN005+LPS). Data are mean ± s.d.

**Supplementary Fig. 4.**
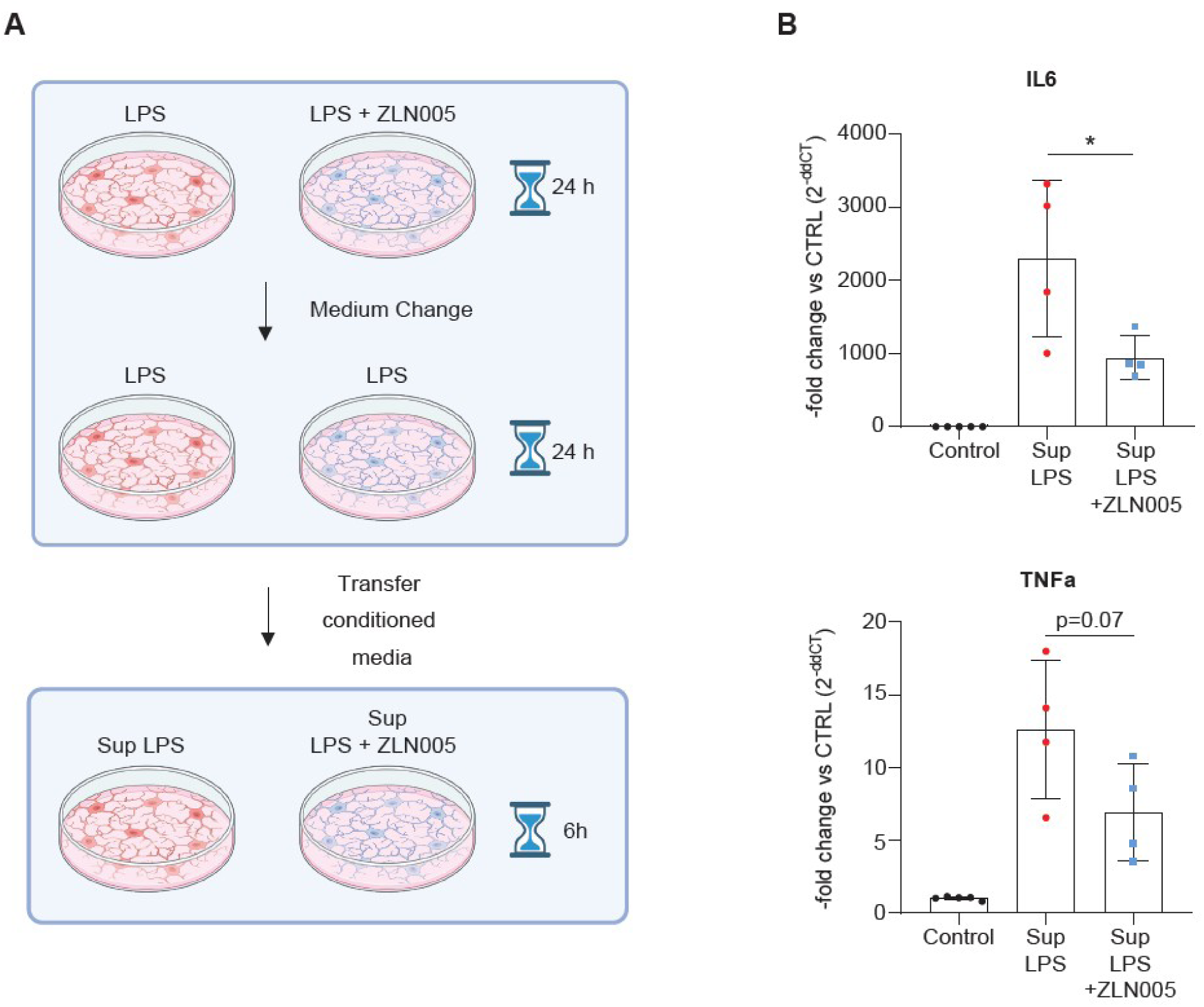
ZLN005 pre-conditioning reshapes the microglial secretome toward a less inflammatory paracrine signal. (**A**) Two-stage conditioned-medium transfer design: donor iMG were treated with LPS or LPS+ZLN005, thoroughly washed to remove ZLN005 and re-challenged with LPS alone, and their supernatants transferred to naive recipient iMG from an independent differentiation. (**B**) qPCR in recipient iMG showing reduced *IL6* induction (*P* < 0.05) and a trend toward reduced *TNFA* (*P* = 0.07) in response to the ZLN005-conditioned secretome relative to the LPS-only secretome. Expression is 2^−ΔΔCt^ fold-change versus untreated control; data are mean ± s.d.

**Supplementary Fig. 5.**
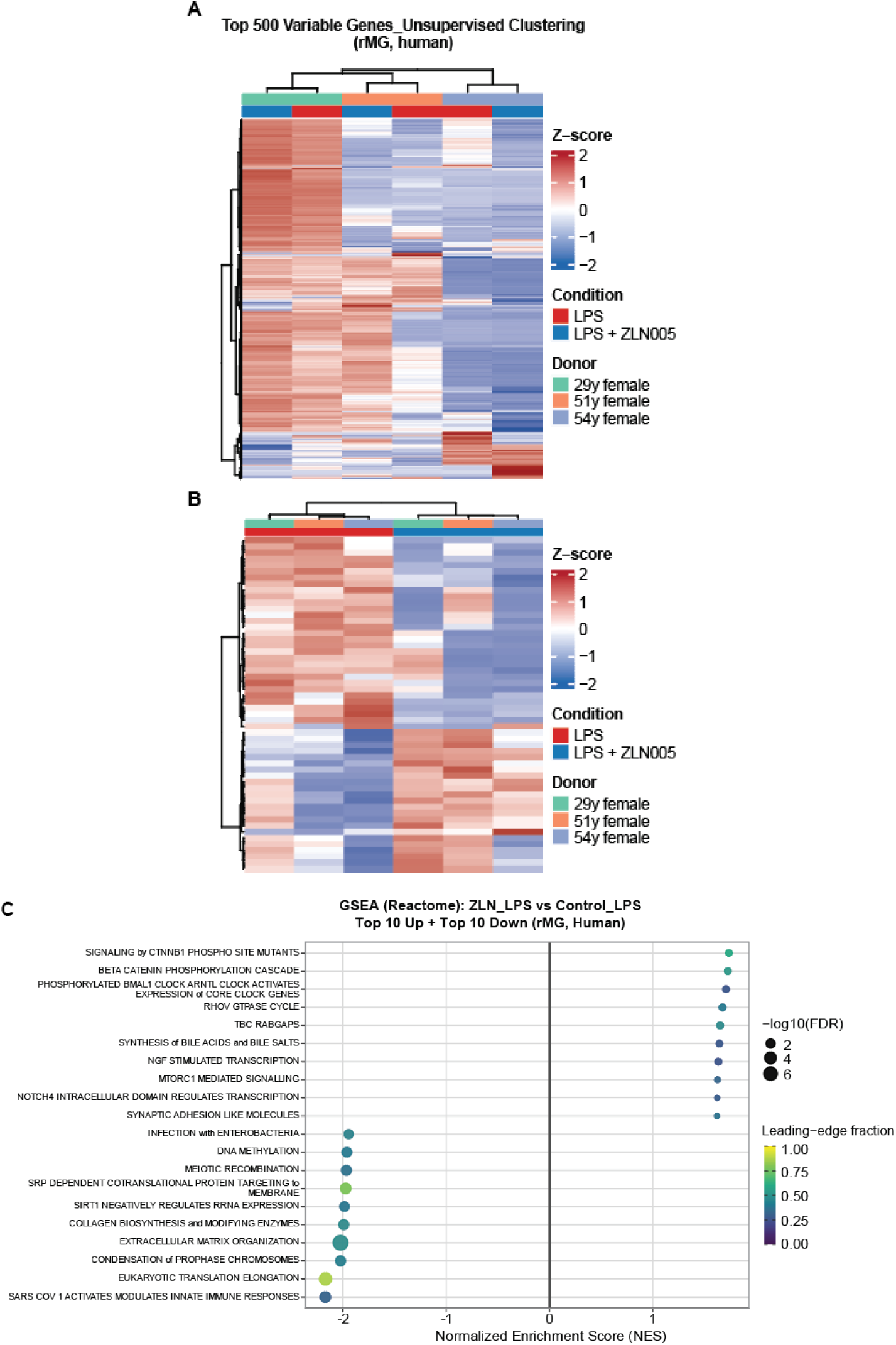
Transcriptional clustering and pathway enrichment in primary human donor retinal microglia (rMG). (**A**) Unsupervised and (**B**) supervised clustering heatmaps of the top 500 most variable genes (row Z-scored), annotated by condition (LPS, LPS+ZLN005) and donor (29, 51, 54 years, female). (**C**) Reactome GSEA for the LPS+ZLN005 versus LPS contrast (top 10 up-and top 10 downregulated pathways). ZLN005 dampens innate-immune, protein-synthesis and extracellular-matrix modules, while several anabolic/growth-associated pathways trend toward induction. Dot size, – log10(FDR); color, leading-edge fraction.

**Supplementary Fig. 6.**
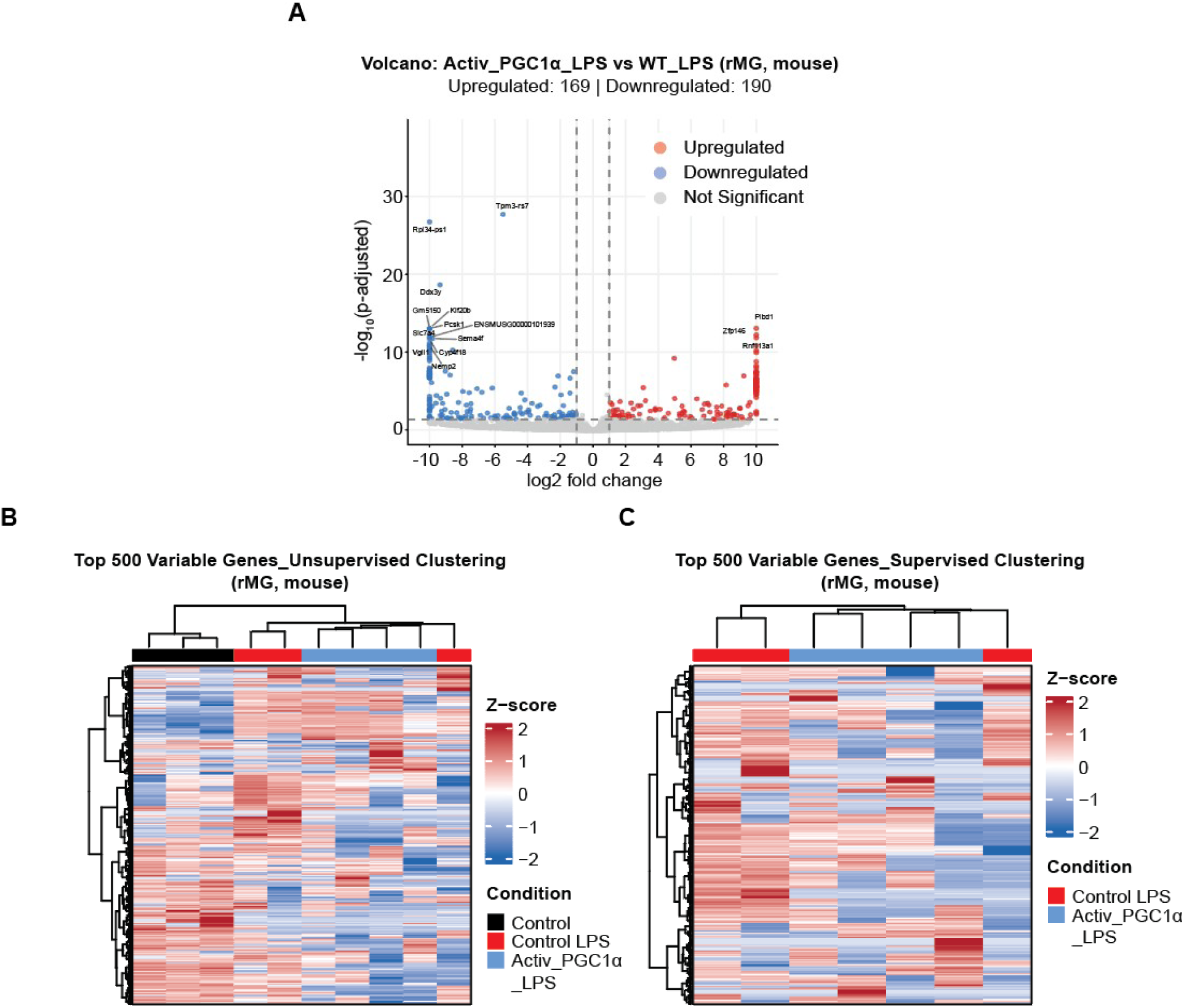
Differential expression and clustering in primary mouse retinal microglia (rMG). (**A**) Volcano plot of the Activ_PGC1α+LPS versus wildtype+LPS contrast (169 upregulated, 190 downregulated genes; DESeq2 Wald test, adjusted *P* < 0.05). (**B**, **C**) Heatmaps of the top 500 most variable genes (row Z-scored): (**B**) unsupervised clustering (control, control+LPS, Activ_PGC1α+LPS) and (**C**) supervised clustering (control+LPS versus Activ_PGC1α+LPS).

**Supplementary Fig. 7.**
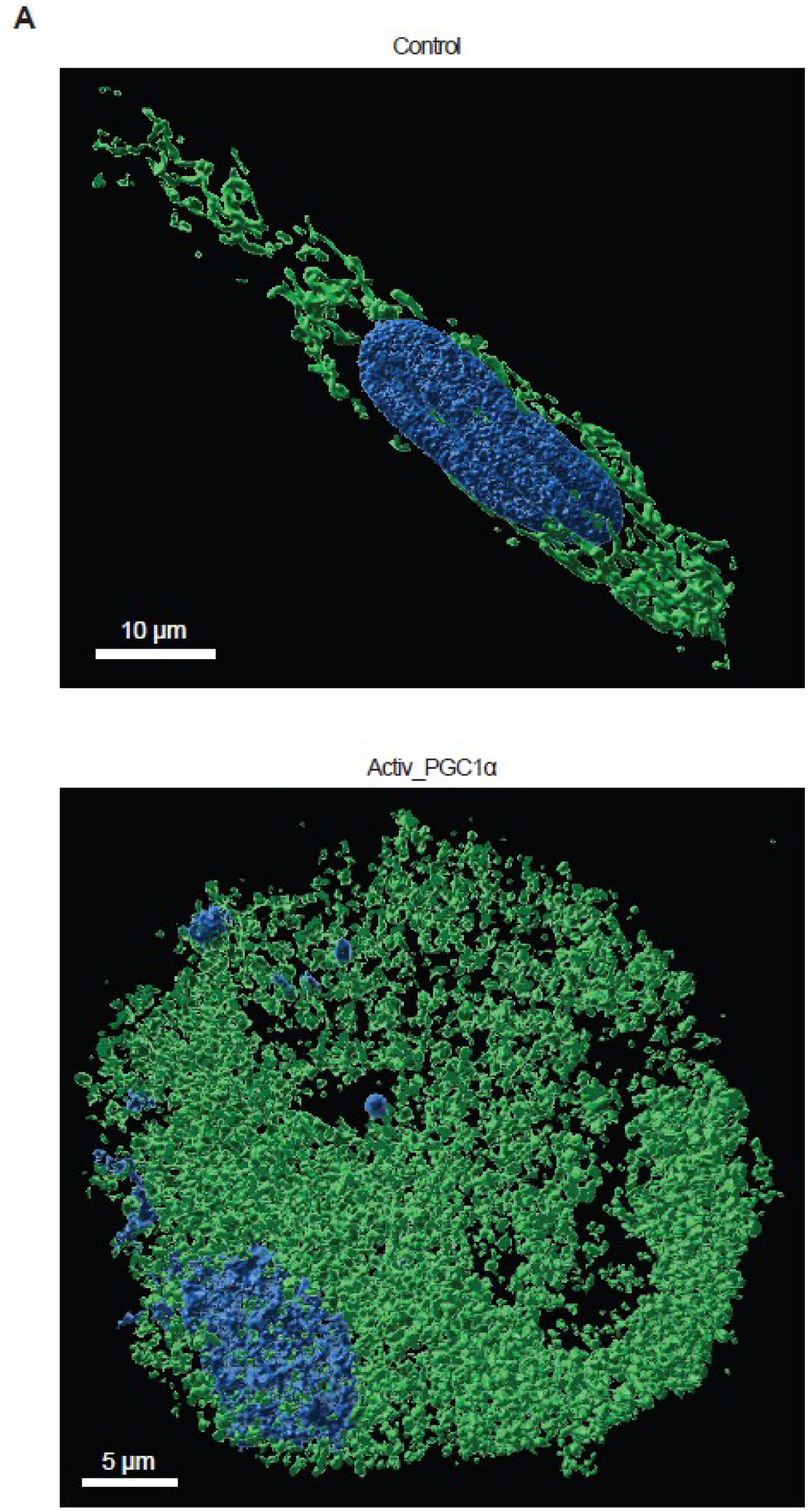
Activ_PGC1α expanded the mitochondrial network in primary retinal microglia. (**A**) Qualitative Airyscan super-resolution imaging of MitoTracker-labeled primary retinal microglia. *Activ_PGC1α*-expressing cells display an aberrant, denser mitochondrial network compared with those of wild-type controls, suggesting organelle-level confirmation of the induced mitochondrial-biogenesis program. Scale bars indicated.

**Supplementary Fig. 8.**
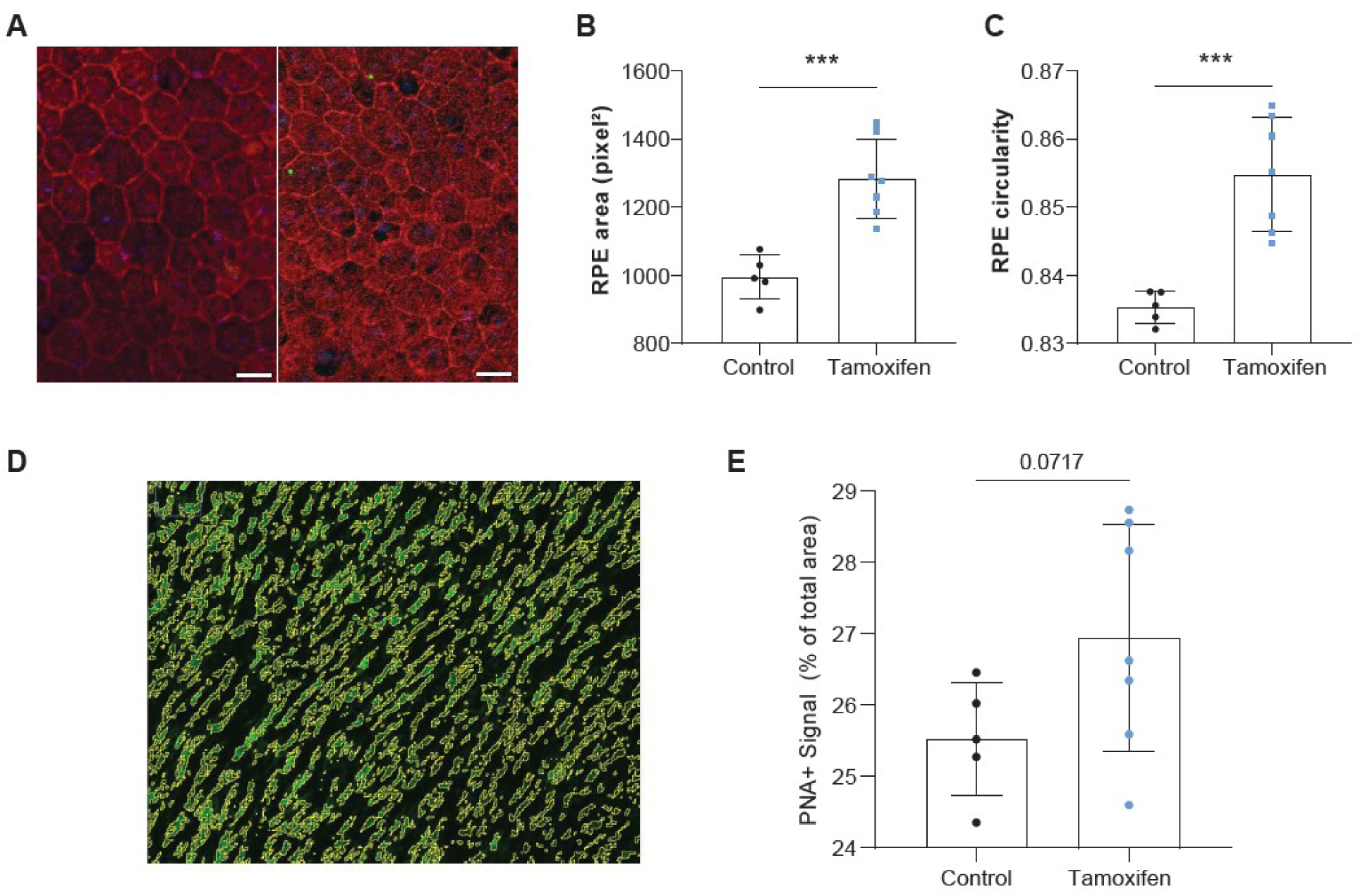
RPE and cone preservation in *Activ_PGC1α*-expressing hu*RHO^P347L/+^*retinae. (**A**) Representative RPE flatmounts stained for the tight-junction marker ZO-1 (tamoxifen - left versus control - right). Scale bar 20 µm. (**B**) RPE cell area and (**C**) RPE circularity, both significantly higher/more uniform in tamoxifen-injected animals (*P* < 0.001), indicating preservation of the regular hexagonal monolayer. (**D**) Representative PNA-labeled cone segmentation and (**E**) PNA+ cone-occupied area (% of total), showing a near-significant trend toward greater cone coverage in tamoxifen-injected animals (Welch, P = 0.072). Data are mean ± s.d

**Supplementary Fig. 9.**
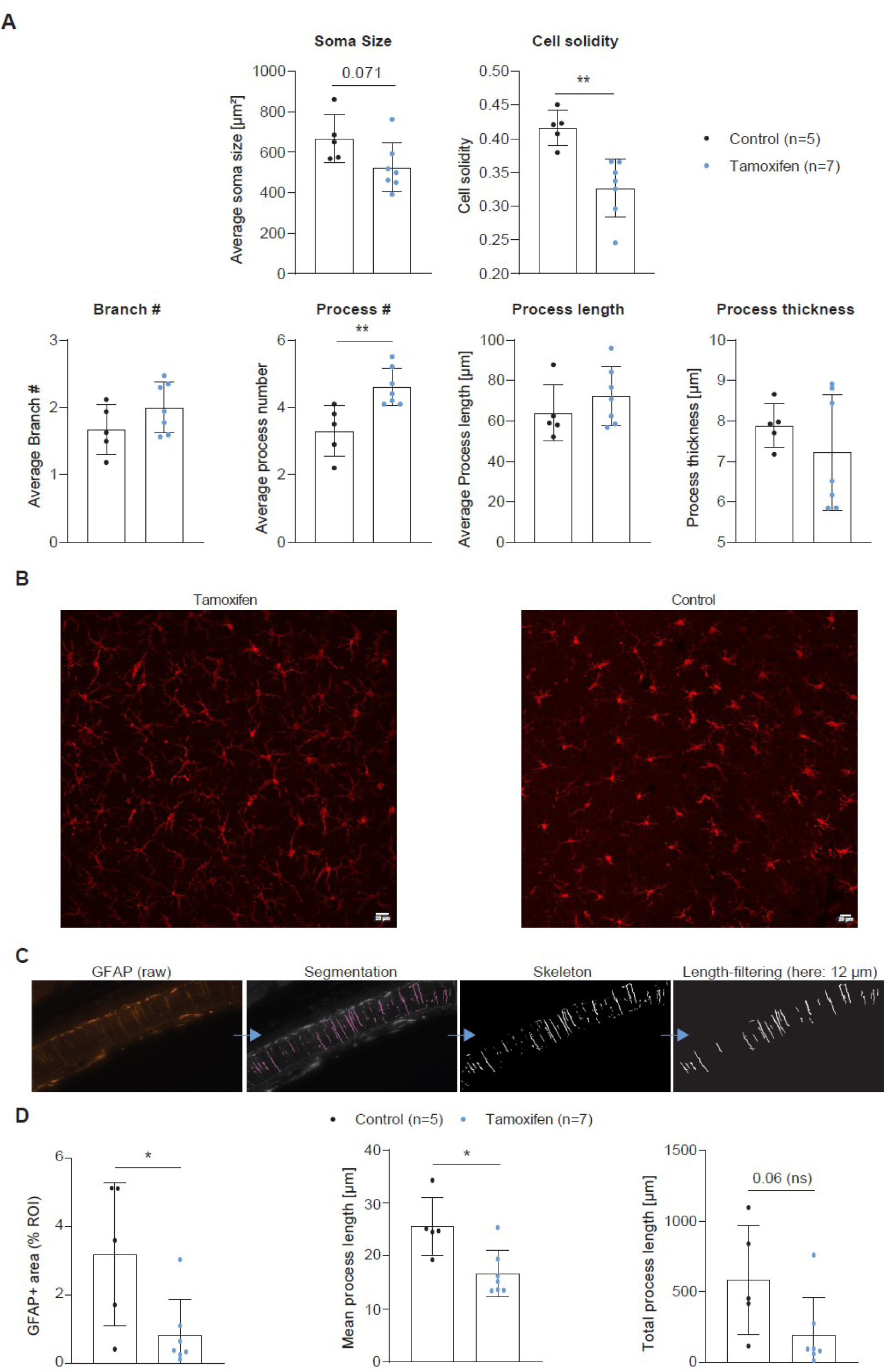
Microglial morphometry and Müller-glia reactivity in *Activ_PGC1α*-expressing hu*RHO^P347L/+^* retinae. (**A**) Quantitative morphometry of IBA1^+^ subretinal cells (control n = 5, tamoxifen n = 7): tamoxifen-injected animals show reduced cell solidity (*P* < 0.01) and an increased number of processes per cell (*P* < 0.01), consistent with a shift from amoeboid toward ramified, homeostatic morphology; soma size, branch number, process length and process thickness do not differ significantly. (**B**) Representative IBA1-stained retinal flatmounts (tamoxifen versus control). Scale bar, 20 µm. (**C**) Image-analysis workflow for GFAP+ Müller-glial process quantification in retinal cross-sections: raw GFAP signal, automated segmentation, application of the ≥12 µm Feret length filter, and the final filtered objects used for measurement. (**D**) Quantification of GFAP^+^ area (percentage of ROI), mean process length, and total process length at the 12 µm Feret cut-off (tamoxifen n = 7, control n = 5). Bars show mean ± SD with individual animal values overlaid. GFAP^+^ area (P = 0.026) and mean process length (P = 0.010) are significantly reduced in tamoxifen animals; total process length shows a non-significant reduction (P = 0.064). Unpaired two-tailed Student’s t-test; *P < 0.05, ns = not significant.

**Supplementary Fig. 10.**
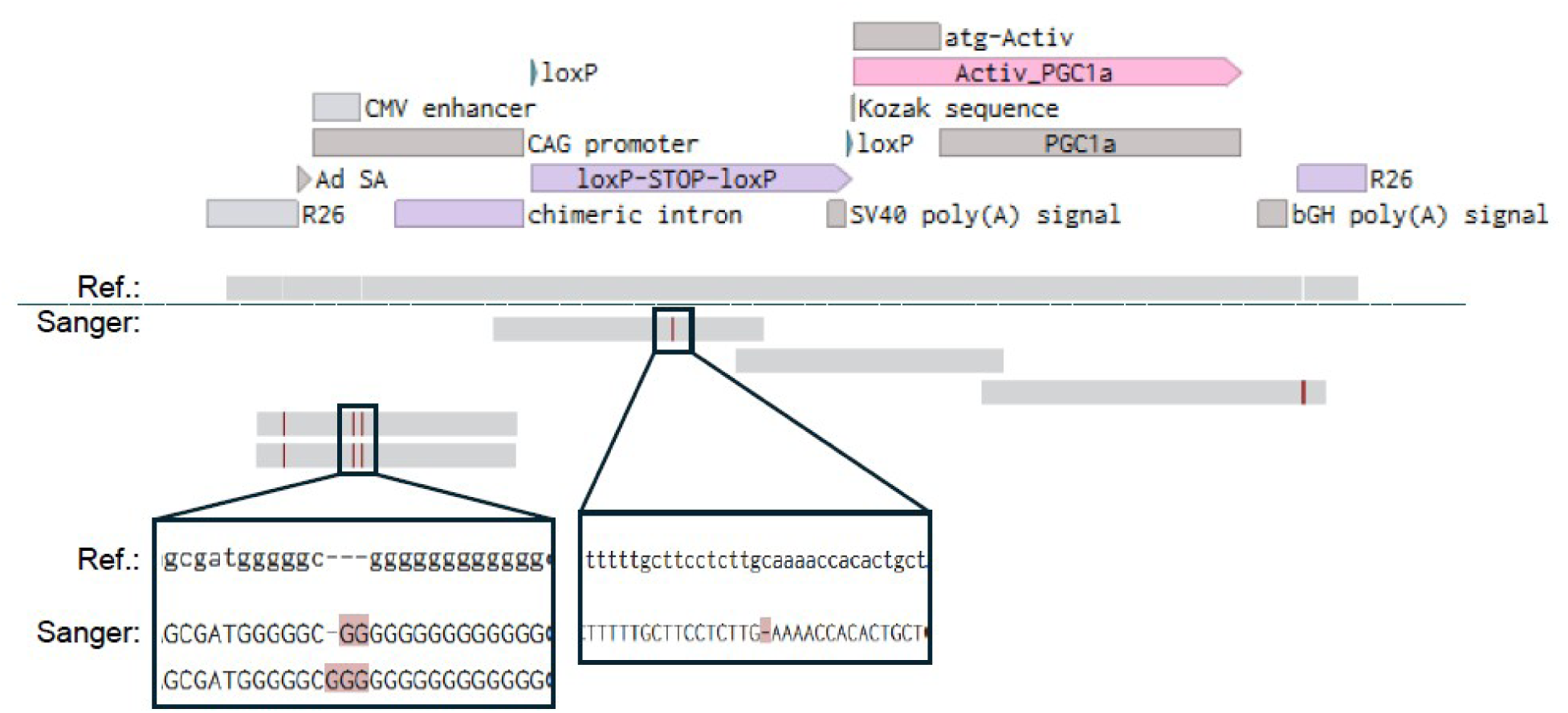
Sanger-sequencing validation of the *R26^LSL-Activ_PGC1α^* knock-in cassette. (**Top**) Annotated map of the targeted *Rosa26* allele (5′→3′): *R26* homology arm, CAG promoter (CMV enhancer, chimeric intron), loxP-flanked transcriptional STOP cassette, the *Activ_PGC1α* transgene (atg-Activ start, Kozak sequence, Activ_PGC1α–*Ppargc1a* ORF), SV40 and bGH poly(A) signals, and the distal *R26* arm. (**Middle**) Coverage of five overlapping Sanger reads tiled across the full cassette, aligned to the *in silico* targeted-allele reference. (**Bottom**) Alignment detail at the two positions that differ from the reference (red): a poly-G run in the CAG-promoter region and an A/T-rich indel within the loxP-STOP-loxP cassette. Both are Sanger homopolymer basecalling artifacts (G-tract compression and A/T-run length ambiguity), and both lie outside the Activ_PGC1α coding sequence, which matched the reference at 100% identity with no discordant bases. Reference (Ref.) and Sanger consensus (Sanger) tracks are indicated at left. Alignment was performed with Auto-MAFFT on Benchling.

**Supplementary Table 1:** Postmortem eye donor information.

| **Donor** | **Age** | **Gender** | **Cause of death** | **Ocular History** | **Relevant Medical History** |
| --- | --- | --- | --- | --- | --- |
| A | 29 y | Female | Liver failure | Cotton wool spots (OU), scleral icterus (OU) | Type 2 diabetes mellitus, systemic lupus erythematosus |
| B | 51 y | Female | Acute respiratory distress syndrome (ARDS) | Keratopathy, transient eyelid edema | Obesity |
| C | 54 y | Female | Subarachnoid hemorrhage | None documented. | Hypertension, history of tobacco use |

**Supplementary Table 2:**
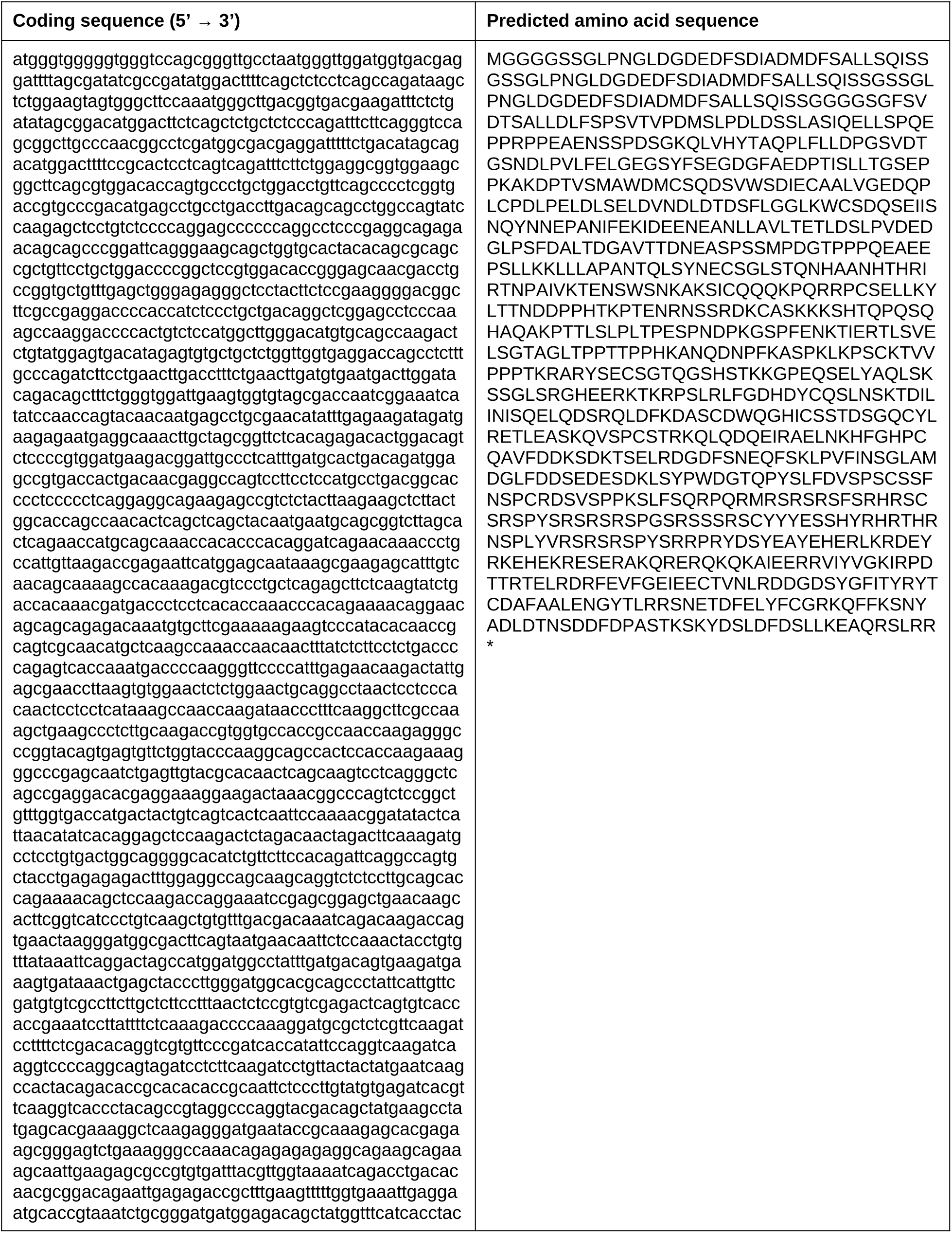
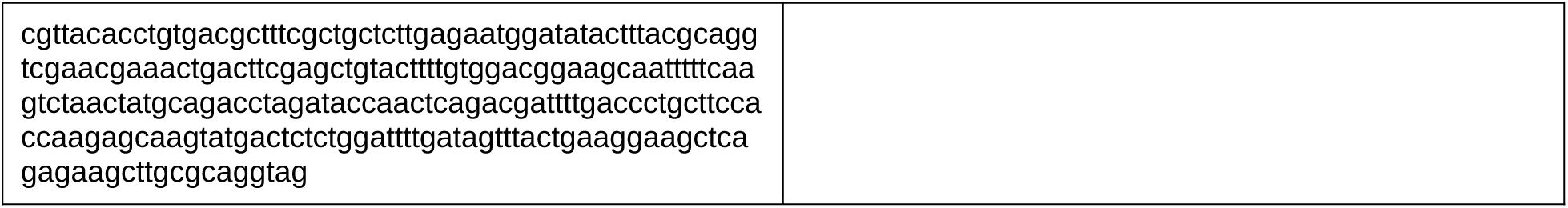
Activ_PGC1α coding sequence (left) and predicted amino acid sequence (right)

**Supplementary Table 3:**
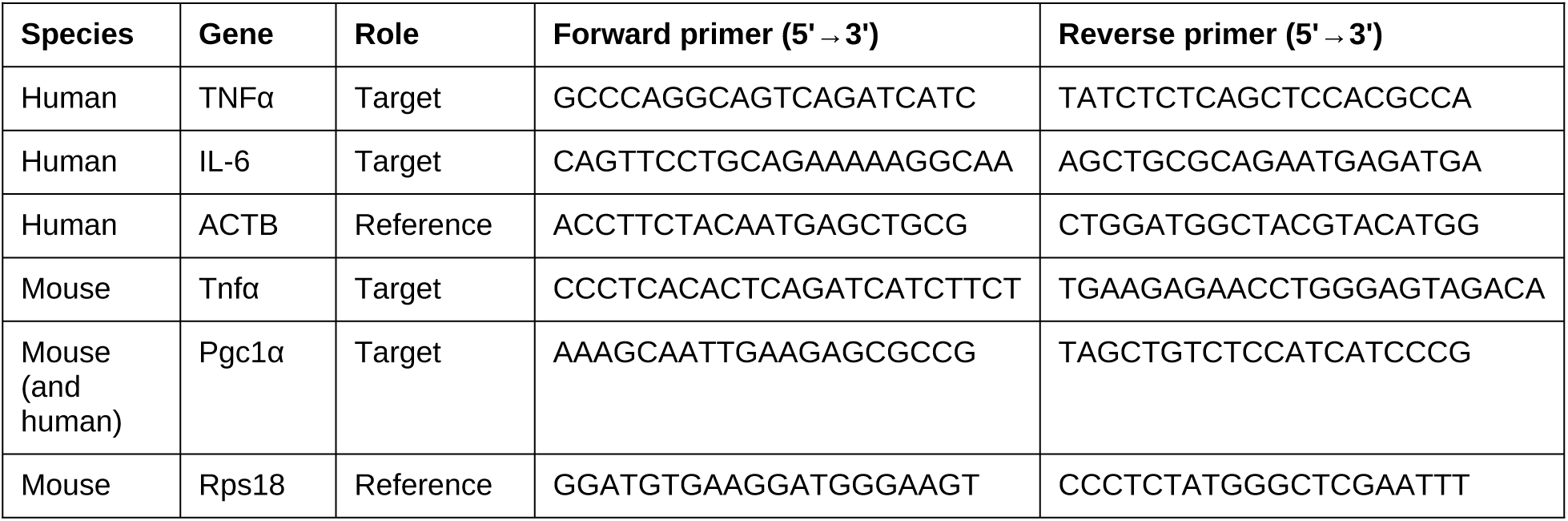
Quantitative PCR primers.

